# When does post-ingestive dopamine signaling occur in humans?

**DOI:** 10.64898/2026.09.25.754497

**Authors:** Valerie L. Darcey, Juen Guo, Kevin D. Hall

## Abstract

The post-ingestive dopamine response is an integral determinant of subsequent eating behavior in animals yet measurement of the post-ingestive response to nutrients in humans has been elusive when assessed in the early post-ingestive period (30 minutes postprandially). We conducted secondary analysis comparing dopamine D2 receptor binding potential (D2BP) measured using [^18^F]fallypride positron emission tomography in two inpatient controlled feeding cohorts consuming standardized diet (50% carbohydrate; 35% fat; 15% protein) with similar characteristics and procedures but differing in time since last meal: confirmed overnight fast (n=57; 66.7% female; BMI 29.6±7.6 kg/m^2^; 31.9±7.2 years old) and 2-hours after a standardized meal reflecting 20% of energy requirements (n=43; 48.8% female; BMI 28.8±7.8 kg/m^2^, 31.3±7.7 years old). Compared to the fasted cohort, the 2-hour postprandial cohort exhibited significantly reduced whole striatal D2BP (p<0.0001) as well as in clusters in insular cortex, prefrontal cortex which remained after adjustment for age, sex, and insulin sensitivity. Voxelwise comparisons of the relationship between D2BP and BMI in the fasted and fed cohorts reveals clusters within dorsolateral and ventral striatum where dopamine level is dependent on postprandial state and adiposity. Given the similarities between protocol procedures and participant characteristics, our results suggest that the post-ingestive dopamine response to nutrients may be prominent in the extended post-ingestive period and warrant investigation within-subject.

## Introduction

Dopamine responses to stimuli play a central role in shaping behaviors including eating. Preclinical evidence suggests that consciously perceived taste and orosensory signals – neither necessary or sufficient to impact food choice long term, e.g. (Ferreira, Tellez et al. 2012) – may function largely as proxy rewards in that they cue the arrival of metabolizable nutrients and accompanying post-ingestive signals that ultimately reinforce food-seeking (de Araujo, Schatzker et al. 2020). These post-ingestive signals conveyed through gut–brain pathways for dietary fats and carbohydrates (DiFeliceantonio and Small 2019) are considered the primary reinforcement signals thought to critically shape food choice (Weber, Yee et al. 2025).

However, efforts to detect a robust post-ingestive dopamine response using positron emission tomography (PET) in humans have yielded inconsistent results. In the largest human study to date, we recently reported that a palatable ultra-processed milkshake produced a highly variable post-ingestive dopamine response that was not significantly different than fasting (Darcey, Guo et al. 2025). Results from a more sensitive PET protocol has suggested that milkshake-evoked post-ingestive dopamine release may display a peak around 20 minutes after ingestion (Thanarajah, Backes et al. 2018) raising possibility that human post-ingestive dopamine signals are subtle and difficult to detect with standard methods.

Critically though, nearly all human studies—including our own—have focused on the first 30 minutes after ingestion. Animal studies highlight that the timing of post-ingestive dopamine measurements are crucial: a brief intragastric nutritive infusion does not impact dorsal striatal dopamine during the early post-ingestive period in mice (Grove, Hakimi et al. 2025) but can ultimately cause sustained increases striatal dopamine for hours thereafter (McDougle, de Araujo et al. 2024). In humans, the postprandial window spans multiple hours during which nutrients, hormones and metabolic signals change dynamically (Berry, Valdes et al. 2020). These considerations raise the possibility that human post-ingestive dopamine responses may emerge later in the postprandial period than typically examined.

Here, we tested whether binding of a dopamine D2 receptor antagonist ([^18^F]fallypride) using positron emission tomography (PET) measured during an extended postprandial window (2–5 hours after a standardized meal) differs from binding measured in the overnight fasted state. In contrast to the highly variable and nonsignificant early postprandial dopamine responses we previously observed (Darcey, Guo et al. 2025), we found significantly reduced receptor availability in the 2-hour post-ingestive state—consistent with increased dopamine release during this extended post-ingestive period.

## Results

Descriptions of the individual pre-registered inpatient clinical trials have been described elsewhere (Guo, Simmons et al. 2014, Darcey, Guo et al. 2025). In brief, 57 weight-stable adults (67.7% female) completed morning [^18^F]fallypride scans in the confirmed overnight fasted (15 hours) state (“Fasted”) while 43 weight-stable adults (48.8% female) completed [^18^F]fallypride scans starting two hours after a standardized breakfast (0800 hours) comprising 20% of energy needs as estimated using indirect calorimetry (“Fed”). A comparison of participant characteristics indicates overlap in demographics, adiposity, and eating behavior traits (**Table 1**). Protocols were similar in main methodological details (**Table 2**). Both cohorts consumed eucaloric standardized diets to stabilize nutrient input (50% of calories from carbohydrate, 35% from fat, 15% from protein; see **METHODS**) provided by the NIH Metabolic Kitchen in advance of PET scanning. At the time of their [18F]fallypride PET scans, participants in the Fasted cohort consumed the standardized diet for a mean of 6.5 days (range 4-8 days) while the Fed cohort consumed the standardized diet for a mean of 4.8 days (range, 4-7 days).

**Table 1.** Comparison of participant characteristics of “Fed” and “Fasted” Cohorts.

|  | **"Fed" Cohort** | **"Fasted" Cohort** | ***p*** |
| --- | --- | --- | --- |
| **N** | 43 | 57 |  |
| **Female** | 21(48.8%) | 38(66.7%) | 0.1 |
| **Age (years)** |  |  |  |
| *Mean* | 31.3±7.7 | 31.9±7.2 | 0.69 |
| *Range* | 20.1-45.7 | 19.0-46.0 |  |
| **Race** |  |  |  |
| *Black* | 26(60.5%) | 28(49.1%) |  |
| *White* | 11(25.6%) | 19(33.3%) |  |
| *Asian* | 2(4.7%) | 7(12.3%) |  |
| *Other* | 4(9.3%) | 3(5.3%) | 0.37 |
| **Body weight (kg)** |  |  |  |
| *Mean* | 86.0±24.6 | 84.6±24.2 | 0.77 |
| *Range* | 44.1-134 | 45.9-149 |  |
| **Body fat (%)** |  |  |  |
| *Mean* | 31.0±11.8 | 34.7±12.1 | 0.14 |
| *Range* | 9.5-51.5 | 11.3-56.7 |  |
| **BMI (kg/m<sup>2</sup>)** |  |  |  |
| *Mean* | 28.8±7.8 | 29.6±7.6 | 0.60 |
| *Range* | 17.8-44.6 | 20.3-44.4 |  |
| **REE (Kcal/d)** |  |  |  |
| *Mean* | 1632±376 | 1614±324 | 0.80 |
| *Range* | 915-2421 | 963-2511 |  |
| **HOMA-IR** |  |  |  |
| *Mean* | 2.2±1.5 | 3.4±3.5 | 0.06 |
| *Range* | 0.7-5.9 | 0.6-22.8 |  |

**Table 2.**
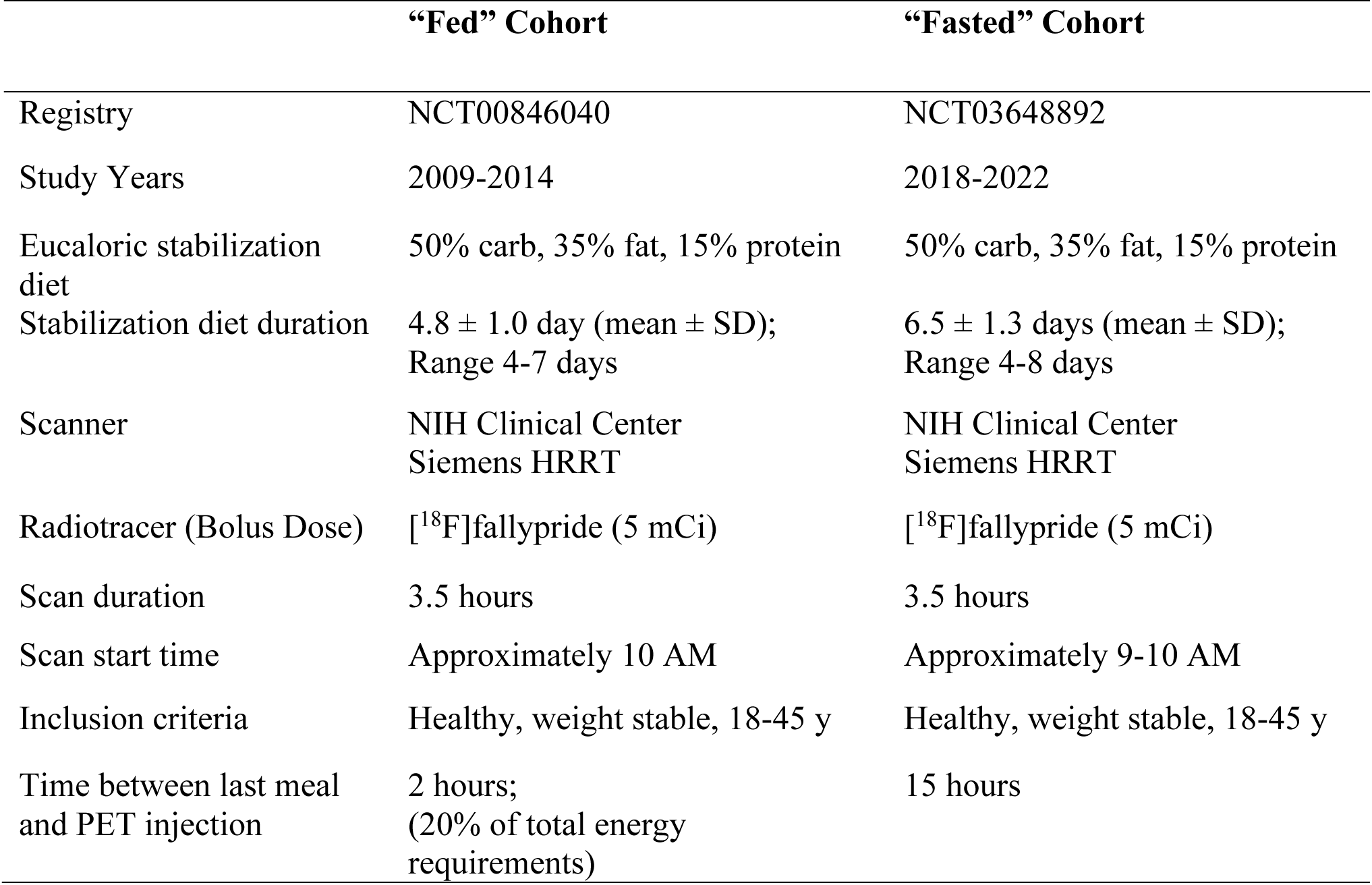
Protocol methods comparison.

|  | **“Fed” Cohort** | **“Fasted” Cohort** |
| --- | --- | --- |
| Registry | NCT00846040 | NCT03648892 |
| Study Years | 2009-2014 | 2018-2022 |
| Eucaloric stabilization diet | 50% carb, 35% fat, 15% protein | 50% carb, 35% fat, 15% protein |
| Stabilization diet duration | 4.8 ± 1.0 day (mean ± SD);<br>Range 4-7 days | 6.5 ± 1.3 days (mean ± SD);<br>Range 4-8 days |
| Scanner | NIH Clinical Center<br>Siemens HRRT | NIH Clinical Center<br>Siemens HRRT |
| Radiotracer (Bolus Dose) | [<sup>18</sup>F]fallypride (5 mCi) | [<sup>18</sup>F]fallypride (5 mCi) |
| Scan duration | 3.5 hours | 3.5 hours |
| Scan start time | Approximately 10 AM | Approximately 9-10 AM |
| Inclusion criteria | Healthy, weight stable, 18-45 y | Healthy, weight stable, 18-45 y |
| Time between last meal and PET injection | 2 hours;<br>(20% of total energy requirements) | 15 hours |

### D2BP was significantly reduced in the Fed Cohort compared to the Fasted Cohort

Participants completed [^18^F]fallypride PET scans in the morning after an overnight in the NIH Clinical Center. The Fasted cohort completed their scans in the overnight fasted state while the Fed cohort initiated their PET scans approximately two hours after a standardized breakfast reflecting 20% of their resting energy requirements as determined by indirect calorimetry (183-484 kcal; 15% kcal protein, 35% fat, 50% carbohydrate). A region of interest (ROI) comparison of two sample mean [^18^F]fallypride D2BP for the whole striatum indicates that D2BP is significantly reduced in the Fed cohort compared to the fasted cohort (26.47 ± 0.41 SE vs 29.65 ± 0.57 SE; p < 0.0001; **Figure 1A**), and remains significant reduced after adjustment for age, sex, and HOMA-IR (26.61 ± 0.44 SE vs 29.54 ±0.52 SE; p < 0.0001; **Supplementary Figure 1A**). Compared to the Fasted Cohort, mean D2BP was significantly reduced in the Fed cohort across striatal subregions of interest including bilateral caudate, right putamen, left accumbens and left pallidum bilaterally including (p’s<0.025; **Figure 1B-E**) and again remained significantly reduced after adjustment for age, sex and HOMA-IR (p’s< 0.05) except for left putamen, right accumbens, and right pallidum (p’s > 0.29 **Supplementary Figure 1B-E**).

**Figure 1.**
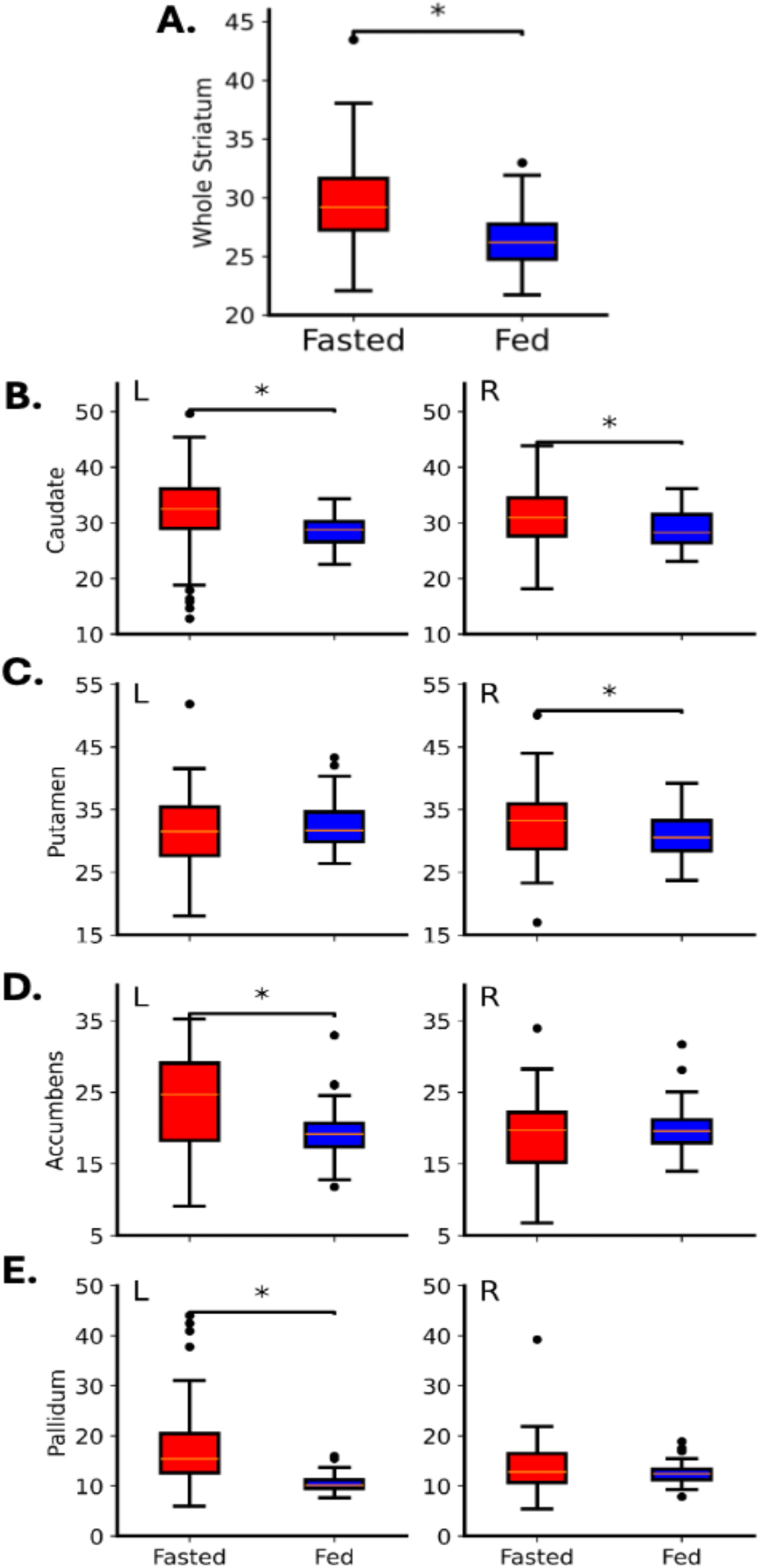
Region of interest dopamine D2R binding potential (D2BP) measured by [^18^F]fallypride in participants the overnight fasted vs. 2-hour post-ingestive state. D2BP (y-axis) is significantly reduced in individuals in the 2-h post-ingestive state (Fed, n=43; blue bars) compared to individuals in the overnight fasted state (Fasted n=57; red bars) across (A) whole striatum Δ BP −3.18 ± 0.75 (p<0.0001), (B) caudate (left, Δ BP 2.87 ± 1.23, p=0.011; right Δ BP −2.29 ± 0.95 p=0.011) (C) putamen (left Δ BP 1.04 ± 1.02, p=0.28; right Δ BP −2.22 ± 1.04, p=0.025), (D) accumbens (left Δ BP −4.2 ± 1.23 p=0.0003; right Δ BP 0.77 ± 0.97, p=0.398) and (E) pallidum (left Δ BP −6.83 ± 1.32 p<0.0001; right Δ BP −0.95 ± 0.86, p=0.225).

We conducted a voxelwise comparison of D2BP between Fasted and Fed cohorts. Given that the relatively high affinity of [^18^F]fallypride for the D2 receptor facilitates quantification of receptor availability in brain regions with 10-100 times lower receptor density than the striatum (Kessler, Whetsell et al. 1993, Mukherjee, Christian et al. 2002), we expanded the voxelwise analyses to encompass the whole brain (*p-*uncorrected<0.0001; cluster defining threshold 30 voxels; false discovery rate (FDR) corrected<0.001). Results of the voxelwise analyses supported results of the striatal ROI analyses above in that significant clusters indicating reduced D2BP in the Fed compared to the Fasted cohort were present in the bilateral striatum, spanning anteriorly to posteriorly in both the dorsal and ventral striatum. Significant extra-striatal clusters indicating decreases in D2BP in the Fed state were also evident in regions including insula, calcarine gyrus, thalamus and regions within the prefrontal cortex (inferior frontal gyrus and anterior cingulate cortex) (**Table 3**; **Figure 2**). Voxelwise results largely remained after adjustment for age, sex and HOMA-IR, with the exception of the anterior cingulate and left putamen (**Supplementary Table 1; Supplementary Figure 2**). There were no clusters which emerged for the reverse contrast (D2BP Fed > D2BP Fasted) in either the unadjusted or adjusted conditions.

**Figure 2.**
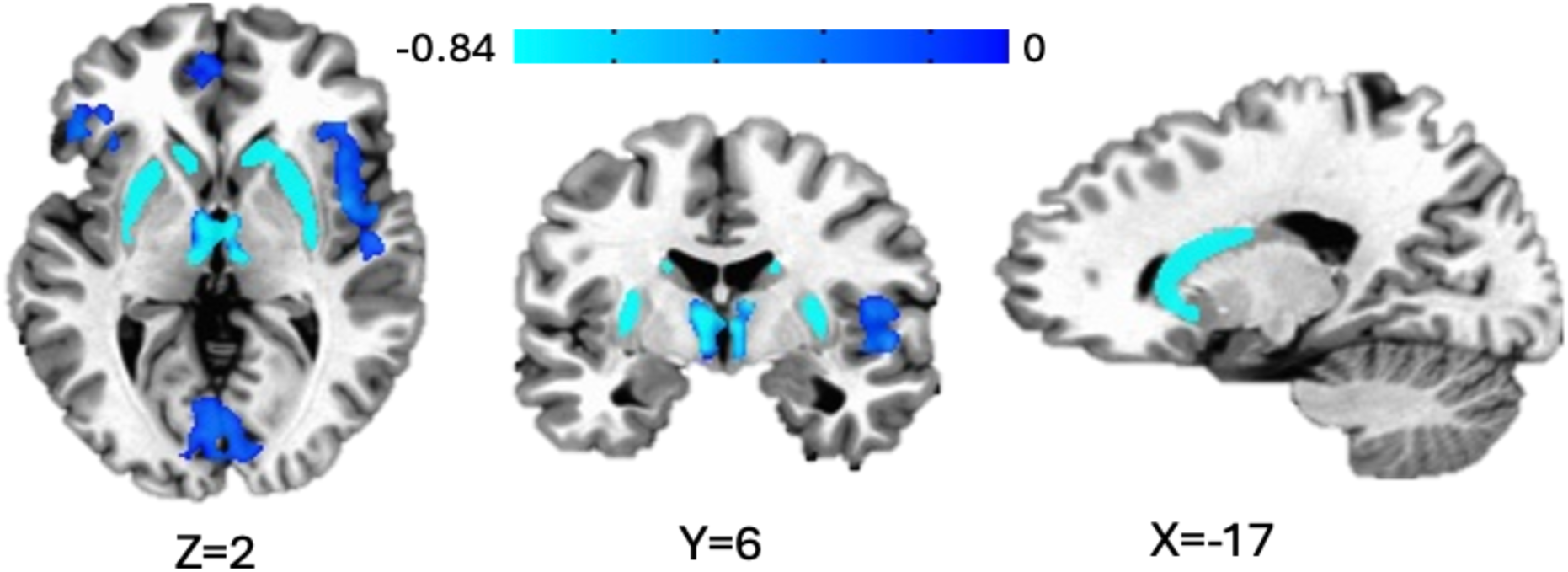
Voxel-wise comparisons of striatal dopamine D2R binding in fed vs. fasting states. Voxel-wise analyses identified clusters across the whole brain where dopamine D2R binding potentials significantly decreased in fed state compared to fasting state. Clusters are defined by faces-touching neighbor clustering, bi-sided uncorrected p < 0.0001, cluster size > 30 voxels, and multiple-comparison corrected p < 0.05. Voxels are displayed after cubic convolution resampling. The color map represents the reduced BP’s in individual voxels (Contrast: D2BP Fed – D2BP Fasted).

**Table 3.** Locations of clusters from voxelwise analyses. PET resolution 3.5mm^3^. Imaging analyses conducted in Analysis of Functional Neuroimaging (AFNI) (clusters defined by voxels with faces touching). Coordinates in Talairach Space.

|  | Location of Peak |  |  | Voxels | Size<br>(mm<sup>3</sup>) | t-stat | alpha |
| --- | --- | --- | --- | --- | --- | --- | --- |
|  | x | y | z |  |  |  |  |
| **Difference Between Cohorts (D2BP Fed – D2BP Fasted)<sup>#</sup>** |  |  |  |  |  |  |  |
| Insula, right | -40.2 | 1 | 3 | 186 | 7975 | -4.77 | <0.001 |
| Inferior frontal gyrus (pars orbitalis), right |  |  |  |  |  |  |  |
| Superior temporal gyrus, right |  |  |  |  |  |  |  |
| Calcarine gyrus, left | 1.8 | 78 | -0.5 | 121 | 5188 | -7.21 | <0.001 |
| Calcarine gyrus, right |  |  |  |  |  |  |  |
| Lingual gyrus, left |  |  |  |  |  |  |  |
| Inferior frontal gyrus (pars triangularis), left | 29.8 | -16.5 | -4 | 117 | 5016 | -4.55 | <0.001 |
| Inferior frontal gyrus (pars orbitalis), left |  |  |  |  |  |  |  |
| Insula, left |  |  |  |  |  |  |  |
| Putamen, right | -29.8 | 8 | 3 | 101 | 4330 | -6.51 | <0.001 |
| Caudate, right |  |  |  |  |  |  |  |
| Pallidum, right |  |  |  |  |  |  |  |
| Thalamus, left | 1.8 | 8 | 6.5 | 58 | 2487 | -6.55 | <0.001 |
| Thalamus, right |  |  |  |  |  |  |  |
| Ventral diencephalon, left |  |  |  |  |  |  |  |
| Anterior cingulate cortex, left | 5.2 | -48 | 3 | 49 | 2101 | -5.89 | <0.001 |
| Middle orbital gyrus, left |  |  |  |  |  |  |  |
| Superior medial gyrus, left |  |  |  |  |  |  |  |
| Putamen, left | 26.2 | 1 | 10 | 41 | 1758 | -7.32 | <0.001 |
| Insula, left |  |  |  |  |  |  |  |
| Caudate, left | 15.8 | -13 | 17 | 40 | 1715 | -4.98 | <0.001 |
| Putamen, left |  |  |  |  |  |  |  |
| **Comparisons of BMI-D2BP slopes ($\beta$ Fed – $\beta$ Fasted)<sup>\*</sup>** |  |  |  |  |  |  |  |
| Putamen, left | 22.8 | -9.5 | 10 | 57 | 2444 | 3.09 | <0.005 |
| Insula, left |  |  |  |  |  |  |  |
| Caudate, left |  |  |  |  |  |  |  |
| Pallidum, left | 29.8 | 18.5 | -0.5 | 57 | 2444 | -1.89 | <0.005 |
| Ventral diencephalon, left |  |  |  |  |  |  |  |
| Putamen, left |  |  |  |  |  |  |  |
| Pallidum, right | -26.2 | 15 | -0.5 | 41 | 1758 | -1.86 | <0.02 |
| Ventral diencephalon, right |  |  |  |  |  |  |  |
| Thalamus, right |  |  |  |  |  |  |  |

*Maximum of top three descriptive labels reflecting largest proportion of cluster provided using the atlas accounting for greatest percent of anatomical localization of cluster.*
*<sup>#</sup> Clusters are defined by faces-touching neighbor clustering, bi-sided uncorrected $p < 0.0001$ , cluster size $> 30$ voxels, and multiple-comparison corrected $p < 0.05$ .*
*<sup>\*</sup> Clusters are defined by faces-touching neighbor clustering, bi-sided uncorrected $p < 0.1$ , cluster size $> 30$ voxels, and multiple-comparison corrected $p < 0.05$ .*

### Fed and Fasted cohorts exhibit distinct voxelwise patterns of correlation between D2BP and BMI

We previously reported that, in a sample with BMI ranging from 18.5-44.4kg/m^2^, D2BP as measured by [^18^F]fallypride in the overnight fasted state was not significantly related to BMI either striatal region of interest or voxelwise analyses (Darcey, Guo et al. 2025). Similarly, we reported that in the extended post ingestive state, mean [^18^F]fallypride D2BP within striatal ROI was also not associated with BMI though we note that the voxelwise correlation patterns suggest the mean binding within the region may be obscuring diametrically opposing binding patterns within anatomical ROIs (Guo, Simmons et al. 2014, Guo, Simmons et al. 2022) (for unthresholded voxelwise binding patterns, see **Supplementary Figure 3A,** Fed; **Supplementary Figure 3B,** Fasted; for unthresholded voxelwise D2BP-BMI correlation patterns adjusted for age, sex and HOMA-IR, see **Supplementary Figure 4A,** Fed**; Supplementary Figure 4B** Fasted). We therefore compared the slopes of correlations between D2BP and BMI both within anatomically defined ROIs and in voxelwise analyses between the Fasted and Fed cohorts.

ROI analyses confirmed that there were no differences in D2BP-BMI correlations between Fed and Fasted cohorts in the striatum as a whole (p = 0.72 **Supplementary Figure 5A)** or any bilateral striatal sub-region of interest: caudate (left p=0.97; right p=0.94), putamen (left p=0.62; right p=0.52), accumbens (left p=0.66; right p=0.77), ventral pallidum (left p=0.15; right p=0.21) (**Supplementary Figure 5B-E)**, even after adjustment for age, sex, and HOMA-IR: whole striatum (p=0.77); caudate (left p=0.88; right p=0.83), putamen (left p=0.68; right p=0.92), accumbens (left p=0.77; right p=0.98), ventral pallidum (left p=0.41; right p=0.38)( **Supplementary Figure 6A-E**).

Voxelwise analyses contrasting D2BP-BMI correlation slopes (β Fed – β Fasted), however, produced clusters in the anterior/dorsolateral region of the striatum (putamen) where the slope of the D2BP-BMI association in the Fed Cohort was significantly *greater* than that of the Fasted Cohort whereas in bilateral ventromedial striatal clusters (pallidum), the slope of the D2BP-BMI association in the Fed Cohort was significantly *less than* the Fasted Cohort (**Figure 3**; **Table 3**). After adjustment for age, sex, and HOMA-IR, two clusters remain in the bilateral ventral striatum where the slope of the D2BP-BMI association in the Fed Cohort was significantly *greater* than that of the Fasted cohort (**Supplementary Figure 7; Supplementary Table 1**).

**Figure 3.**
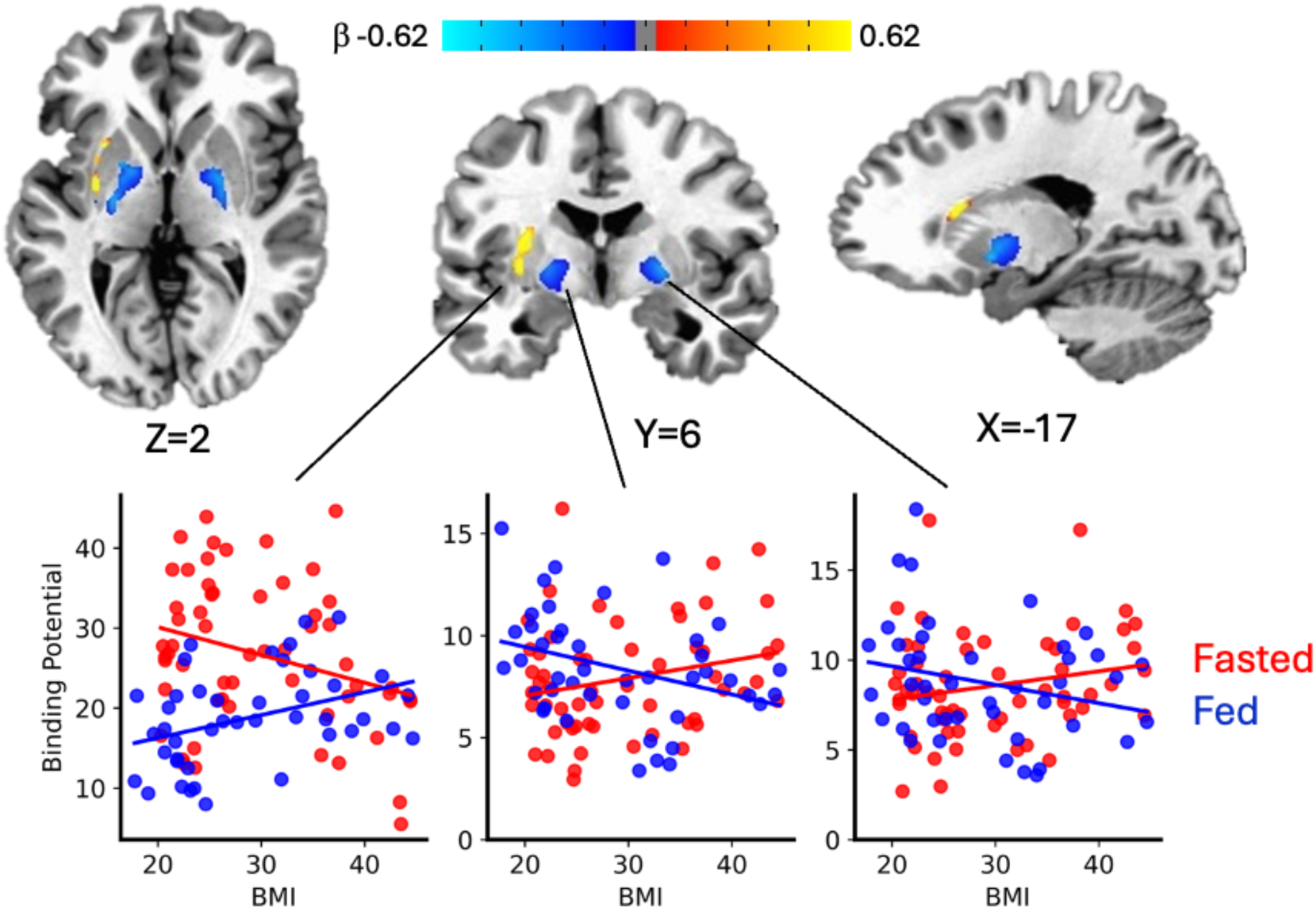
Voxelwise comparison of correlation between dopamine D2 binding potential and adiposity in Fed and Fasted cohorts. Voxel-wise analyses within striatal mask identified clusters where the regression β coefficients of the D2BP-BMI relationship are significantly different in fed vs fasted cohorts in the left dorsolateral putamen (left plot), left pallidum (middle plot) and right pallidum (right plot. Clusters are defined by faces-touching neighbor clustering, bi-sided uncorrected p < 0.1, cluster size > 30 voxels, and multiple-comparison corrected p < 0.05. Voxels are displayed after cubic convolution resampling. The color maps represent the difference of the regression β-coefficients between the fed and fasting states, displayed in cluster-specific scatter plots for visualization purposes.

## Discussion

If post-ingestive dopamine signaling is a central driver of ingestive behavior, an unresolved question is *when* dietary manipulations should be expected to influence dopamine responses in humans. Notably, prior attempts to measure dopamine responses to nutritive stimuli via standard positron emission tomography (PET) scanning approximately 30 minutes after ingestion—including our prior milkshake study the largest of its kind to date (Darcey, Guo et al. 2025)—have yielded largely null and mixed results (Small, Jones-Gotman et al. 2003, Wang, Tomasi et al. 2014, Thanarajah, Backes et al. 2018, Eisenstein, Black et al. 2020, Carnell, Steele et al. 2023). Our null findings have been critiqued on the grounds that dopamine release was assessed outside the relevant temporal window, with the suggestion that measurements occurred *too late* to detect relevant dopamine release (Hall and Darcey 2025). The present data challenge this interpretation. Instead, the results of our exploratory secondary analyses here suggest that prior studies may have attempted to assess dopamine signaling *too early* in the post-ingestive period. We observed a significant reduction in striatal dopamine D2 receptor binding potential in participants scanned in the period spanning 2 to 5 hours after consumption of a standardized meal relative to individuals scanned in the overnight fasted state. We further find exploratory evidence suggesting that extended post-ingestive dopamine release in isolated areas within the striatum may be dampened at higher levels of adiposity. While these findings warrant empirical confirmation, together these patterns intriguingly suggest that endogenous dopamine response to nutrients in the gut occurs on a longer timeline in humans than implicitly assumed by previous neuroimaging work – in the *extended* post-ingestive period.

The delayed dopamine effects we observed in humans may help reconcile the apparent discrepancy between strong preclinical evidence for post-ingestive dopamine signaling and repeated inconsistent and most recently null findings in human PET studies cited above. Rodent studies demonstrate that intragastric nutrient infusions cause striatal dopamine response that scales with the caloric load (Ferreira, Tellez et al. 2012). Energy restriction from fasting decreases striatal dopamine tone (increased D2R availability) in rodents relative to ad libitum–fed controls (Thanos, Michaelides et al., 2008). Our observation of comparatively reduced synaptic dopamine measured in the fasted cohort is therefore consistent with the observation of greater striatal D2 receptor availability in prior preclinical studies. Fasting is known to enhance motivational drive and potentiate phasic dopaminergic responsiveness to subsequent stimuli (Carr 2007). Despite reports that spontaneous dopaminergic activity - and thus tonic dopamine levels - appear to be augmented during fasting potentially mediated by ghrelin (Abizaid, Liu et al. 2006, Cone, McCutcheon et al. 2014, van der Plasse, van Zessen et al. 2015), the magnitude of dopamine release observed here in response to extended postprandial signals (approximately 2–5 hours postprandially) is significantly greater than the net result of tonic dopamine activity in the fasted state.

These findings suggest that conventional post-meal PET protocols may systematically underestimate post-ingestive dopamine responses. Prior human PET studies of post-ingestive dopamine signaling have largely used within-subject designs using [¹¹C]raclopride, a D2 receptor antagonist with high sensitivity to displacement by endogenous dopamine, under conditions expected to maximize detection of effects beginning 10-30 minutes after ingestion (Small, Jones-Gotman et al. 2003, Wang, Tomasi et al. 2014, Thanarajah, Backes et al. 2018, Eisenstein, Black et al. 2020, Carnell, Steele et al. 2023, Darcey, Guo et al. 2025). Despite these favorable design features, these studies—including our own—largely failed to detect robust dopamine responses. The variability and inconsistency across prior studies, together with our null findings following a UPF milkshake at 30 minutes, suggest that early post-ingestive dopamine responses in humans are modest and difficult to detect reliably. In contrast, the present between-subjects comparison using [¹⁸F]fallypride—a tracer with comparatively lower sensitivity to endogenous dopamine displacement—demonstrates a post-ingestive reduction in D2BP when scans are acquired later in the postprandial period. It is important to note the possibility that, in addition to the differences in timing, differences in meal form and composition may have contributed to the differences in binding observed between the present analysis and our prior null investigation. The Fed cohort in the present analysis of the delayed post-ingestive dopamine response consumed a solid meal providing 183–484 kcal (∼20% of resting energy requirements; 15% protein, 35% fat, 50% carbohydrate), whereas the prior study investigating early post-ingestive dopamine response used a 418-kcal milkshake (∼26.7 ± 5.3% of resting energy requirements; 7% protein, 60% fat, 33% carbohydrate). However, integrating findings across tracers and time points supports the interpretation that post-ingestive dopamine signaling in humans emerges more robustly at later time scales, yielding effects of greater magnitude that are more readily detectable even with less displacement-sensitive ligands.

Our results further provide exploratory evidence suggesting that the dopamine response in the extended post ingestive period may be altered in the presence of obesity. Importantly, we observed differences in the relationship between striatal D2 receptor binding potential and adiposity between cohorts in distinct metabolic states that were evident in voxelwise, but not region-of-interest (ROI), analyses. This dissociation supports the inference that null ROI effects may be due to the cancellation of opposing subregional patterns rather than a true absence of adiposity-related differences. Inspection of unthresholded voxelwise correlation maps revealed a striking general reversal in the direction of the BMI–D2BP relationship between the fasted and delayed post-ingestive states. Notably, this pattern contrasts with our previous within-subject study of *early* post-ingestive responses to milkshake ingestion, in which neither ROI nor voxelwise analyses detected an effect of adiposity (Darcey, Guo et al. 2025). These inconsistences underscore the importance of considering both temporal phase of digestion and preserving voxelwise spatial information when probing nutrient–dopamine interactions.

In the extended post-ingestive state, voxelwise analyses revealed that higher BMI was associated with relatively greater dopamine levels in bilateral ventral striatal regions (pallidum), alongside attenuated dopamine levels in a more dorsolateral striatal cluster (putamen). The ventral pallidum is a major output structure of the ventral striatum, integrating limbic information and projecting to thalamic targets that drive motivated incentive signals into action (Smith, Tindell et al. 2009, Soares-Cunha and Heinsbroek 2023). It is possible that greater pallidal dopamine levels in the extended post-ingestive state may may bias behavioral responses toward continued motivation for food seeking despite recent intake (O’Doherty, Deichmann et al. 2002, Small, Veldhuizen et al. 2008). In contrast, dorsal striatal dopamine is responsive to gastrointestinal nutrient signals (Ferreira, Tellez et al. 2012, Han, Tellez et al. 2016). It is interesting to speculate that the attenuated dorsolateral striatal dopamine levels observed during the extended post-ingestive state among individuals with obesity may reflect reduced physiological contrast between fasted and fed metabolic states, limiting dopaminergic discrimination of nutrient status. Importantly, this pattern emerged even though the meal consumed by the Fed cohort was standardized to provide 20% of each participant’s individually estimated energy requirements, indicating that proportional caloric loads may be differentially encoded through dopamine signaling as a function of adiposity. The proximal mechanisms for these effects have yet to be well defined but may include the possibility that, in the presence of obesity, elevated tonic dopamine while fasting (Darcey, Guo et al. 2025) attenuates evoked dopamine responses (Ford 2014) from digestion thereby compressing the dynamic range in synaptic dopamine over which post-ingestive signals are encoded. Additionally, any impairments in central signaling in factors such as leptin (Obradovic, Sudar-Milovanovic et al. 2021) or insulin (Heni 2024) could blunt discrimination between fasted and fed metabolic states, and disrupt reinforcement updating following food intake (Farooqi, Bullmore et al. 2007). Although these mechanisms remain speculative and warrant further within-subject testing, the present results lend support for the idea that regional striatal dopaminergic sensitivity to nutritive load may vary with adiposity in humans and influence the role of dopamine in food reinforcement (de Araujo, Ferreira et al. 2012).

Our analyses also suggest that the extended post ingestive state is associated with network-level dopaminergic modulation outside of the striatum. Though both radioligands are D2/3 receptor antagonists, [^11^C]raclopride and [^18^F]fallypride provide overlapping but not identical information about the dopamine system (Darcey, Guo et al. 2025). [¹⁸F]fallypride has markedly higher affinity (Kd ≈ 0.03 vs. 1.2 nM) which enables reliable quantification in regions with low D2/3 receptor density (Kessler, Whetsell et al. 1993, Mukherjee, Christian et al. 2002). Though longer scan time requirements (∼3 h versus ∼1 h for [^11^C]raclopride) presumably limits use in repeat-scan bolus displacement paradigms (Olsson and Farde 2001), [^18^F]fallypride binding remains sensitive to dopamine efflux from evoked by cognitive tasks (Ceccarini, Vrieze et al. 2012, Miederer, Buchholz et al. 2025) and methylphenidate, which reduces striatal binding by 5–11% (Riccardi, Li et al. 2006, Cropley, Innis et al. 2008, Rominger, Wagner et al. 2010). Leveraging this, our whole-brain voxelwise analyses show that, compared to the fasted state, adults in the extended post-ingestive state exhibit elevated dopamine levels (i.e. reduced D2 binding potential) across a distributed thalamocortical network involved in integrating gustatory and visceral interoception (insula (Avery, Kerr et al. 2015), valuation (orbitofrontal cortex/medial prefrontal (Rolls and Grabenhorst 2008)) identifying and prioritizing relevant external or internal stimuli to optimally guide behavior (anterior cingulate, thalamus and salience network(McCutcheon, Nour et al. 2019) and modulating sensory gain and attentional prioritization (visual cortex; (Vickery, Chun et al. 2011, Yousif, Fu et al. 2016) and cognitive/behavioral control (anterior cingulate/inferior frontal gyrus (Holroyd, Nieuwenhuis et al. 2004, Guitart-Masip, Huys et al. 2012). These widespread decreases in D2 receptor binding potential intriguingly suggest that the post-ingestive dopamine response may involve integration of internal physiological state and valuation/motivational salience with modulation of sensory and behavioral control regions.

If post-ingestive signals operating over hours, rather than minutes, are key contributors of gut-based reinforcement signals for shaping food choice and learning, what are the biological stimuli with relevant kinetics and access to affect central dopamine? Dietary carbohydrate and fat engage separable but interacting gut–brain pathways (McDougle, de Araujo et al. 2024) and their downstream metabolic signals are processed on markedly different temporal scales (Berry, Valdes et al. 2020). Carbohydrates are rapidly digested and absorbed via the hepatic portal circulation, producing swift elevations in plasma glucose and insulin. Glucose can directly drive striatal dopamine efflux by serving as an immediate cellular energy substrate (Tellez, Ren et al. 2013), and insulin independently modulates dopamine signaling (Patel, Carr et al. 2023). However, in our prior human PET study, neither glycemic nor insulinemic responses to a milkshake predicted interindividual variability in dopamine release during the early post-ingestive period (Darcey, Guo et al. 2025), arguing against these rapid signals as primary drivers of delayed dopamine effects. In contrast, dietary fat is digested, absorbed, and reassembled into triglyceride-rich lipoproteins that enter the circulation via the lymphatic system, producing delayed and prolonged postprandial dynamics, fluctuating for several hours after meal ingestion (Berry, Valdes et al. 2020, Bozzetto, Della Pepa et al. 2020). Triglycerides and triglyceride-rich particles can cross the blood–brain barrier, where they serve as metabolizable energy substrates and directly influence dopamine signaling and reward processing (Cansell, Castel et al. 2014, Banks, Farr et al. 2018, Berland, Montalban et al. 2020, Kumar, Wu et al. 2025). Meal-induced elevations in plasma triglycerides – which are impacted in obesity (Wilson, Maes et al. 2021)– are associated with altered striatal responses to palatable taste stimuli in humans (Sun, Veldhuizen et al. 2014). While additional intermediate signals may also contribute, it is interesting to speculate that the delayed kinetics, central accessibility, and known dopaminergic effects of triglycerides position postprandial lipid signaling as a biologically plausible mechanism linking nutrient ingestion to dopamine release during the extended post-ingestive period, making them a compelling candidate for further mechanistic investigation.

Finally, these results may inform an updated interpretation of aspects of our prior work. We previously interpreted the observed reduction in [^18^F]fallypride D2BP following 5 days of a reduced-fat diet compared to baseline diet as evidence of increased dopamine *tone* because all scans were conducted without *acute* nutrient stimulation (Darcey, Guo et al. 2023). However, the present findings suggest that the post-ingestive dopamine response may extend well beyond the period previously considered “acute”, and raise the possibility that observed reduction in D2BP in our prior study reflected not only changes from 5 days of reduced-fat diet exposure but *also* to a delayed dopamine response to the reduced fat meal consumed 2 hours prior to scanning. It is important to note that this interpretation is specific to adults with obesity, characterized by increased dopamine tone in the fasted state (Darcey, Guo et al. 2025). Elevated tonic DA would be expected to shape stimulus evoked phasic DA responses (Grace 2000) potentially attenuating stimulus evoked DA release (Sander, Hooker et al. 2013). While we did not observe a blunted dopamine response to milkshake in the early post-ingestive period (Darcey, Guo et al. 2025), the present results suggest adiposity-dependent attenuation may be observed later in the post-ingestive period. Importantly, however, a post-ingestive measurement at a single timepoint may not fully capture the nature or extent of altered dopamine responses in obesity. Differential dopamine responses to specific nutrients (van Galen, Schrantee et al. 2023) may be shaped by obesity-related alterations in postprandial metabolic dynamics beyond response magnitude alone (e.g., time to peak plasma concentrations and duration/clearance (Mamo, Watts et al. 2001, van Vliet, Koh et al. 2020). Thus, post-ingestive dopamine responses may reflect an interaction between not only background diet and stimulus nutrient composition but also adiposity-related differences in the dynamics of postprandial metabolism.

## Limitations

The most likely explanation for the observation of decreased D2BP in participants scanned in an extended post-ingestive state compared to a cohort with similar characteristics scanned in the overnight fasted state is that significant post-ingestive dopamine signaling occurs later in response to a meal. However, the exploratory nature of the secondary analyses, along with between-subject comparisons across metabolic states, warrants cautious interpretation and motivates targeted within-subject follow-up studies. To confirm these results are not discrepant between tracers, future studies should investigate the late post-ingestive period using [^11^C]raclopride. To further facilitate comparison with available literature, future studies should use high fat, high sugar liquid stimuli to confirm that the D2BP differences observed 2h after eating in the present study are driven by properties of the meal (i.e. lower-fat, solid food). Also, both sex and age have been identified as relevant factors in DA signaling (Riccardi, Zald et al. 2006, Dang, Samanez-Larkin et al. 2016) though these cohorts are similar in age and sex.

Metabolic state modulates cerebral blood flow (Tataranni, Gautier et al. 1999, Gautier, Chen et al. 2000) but two considerations argue against the interpretation that the present results are due to perfusion-related effects on tracer delivery or kinetics rather than changes in endogenous dopamine. First, our binding outcome measure is the distribution volume ratio – an in vivo approximation of (*B_max_*/*K_d_*) + 1—is theoretically insensitive to global changes in CBF because it normalizes target region uptake to a reference region (cerebellum)(Logan, Fowler et al. 1996). This ratio cancels first-order effects of tracer delivery provided that relative tracer transport between regions remains stable. Second, our modeling assumes a constant ratio *R = k_2_/k_2_^’^ = k_1_/k_1_^’^* between target and reference regions. Under this assumption, coordinated changes in delivery and washout (e.g., feeding induced vasodilation) should similarly affect both regions, minimally impacting estimates of the distribution volume ratio. While prior simulations have suggested that binding potential estimates may be sensitive to kinetic parameters, these simulations typically vary one parameter in isolation and do not reflect physiological conditions—such as feeding—where both kinetic parameters are likely to change in tandem. Moreover, those simulations rely on plasma input functions, whereas our reference tissue approach does not (Slifstein and Laruelle 2001). Together, these points reduce concern that a “larger hose” (i.e., increased perfusion) in the fed state artifactually lowers D2BP. Finally, our extra-striatal findings should be interpreted cautiously. In the present design, striatal and extra-striatal binding were estimated simultaneously, and [¹⁸F]fallypride uptake and washout kinetics differ substantially across regions as a function of receptor density, with slower kinetics in striatum relative to cortex (Ceccarini, Vrieze et al. 2012). Nonetheless, the high affinity of [¹⁸F]fallypride provides a unique window into delayed, post-ingestive dopamine responses in brain regions that are inaccessible with lower-affinity tracers such as [^11^C]raclopride.

## Conclusion

The present analyses suggest that the internal milieu in the extended post ingestive period may better reflect the neurobiologically relevant basis for learned associations between sensory cues and metabolic consequences. The elevated dopamine observed 2 hours post-ingestion suggests that a distributed network spanning brain regions canonically implicated in valuation (striatal), interoceptive (insula), and control (frontal/thalamic) tracks internal physiological state over extended timescales beyond a transient reward response to food. Future work should employ within-subject designs with multiple post-ingestive time points to directly map the temporal profile of dopamine release following nutrient consumption. Understanding how post-ingestive dopamine dynamics may differ between stimuli varying in nutrient composition in individuals with and without obesity might shed light on how these states influence food cravings and eating behavior.

## Resource Availability & Lead Contact

Request for further information and resources should be directed to and will be fulfilled by the Lead Contact, Valerie Darcey

## Materials Availability

This study did not generate materials.

## Data and Code Availability

Data from consenting subjects will be available upon request from corresponding author.

## Acknowledgments

This research was supported [in part] by the Intramural Research Program of the National Institute of Diabetes and Digestive and Kidney Diseases (NIDDK) within the National Institutes of Health (NIH). The contributions of the NIH author(s) are considered Works of the United States Government. The findings and conclusions presented in this paper are those of the author(s) and do not necessarily reflect the views of the NIH or the U.S. Department of Health and Human Services.

## Author Contributions

V.L.D. and K.D.H. conceived and designed the study. J.G. conducted the statistical analyses. All authors contributed to interpretation of the results, critically revised the manuscript for important intellectual content, and approved the final version.

## Methods

The secondary analyses described herein were approved by the National Institutes of Health (NIH) Institutional Review Board (IRB) (IRB002590). The parent studies were also approved by the NIH IRB (09-DK-0081; 18-DK-0132), and all participants provided written informed consent prior to participation.

### Participants

#### Postprandial Cohort

Forty-three adults provided informed consent to participate in an inpatient study of striatal dopamine type 2 receptor binding potential (ClinicalTrials.gov NCT00846040). Study outcomes were reported elsewhere regarding the primary metabolic (Hall et al., 2015) and primary PET outcomes (Guo, Simmons et al. 2014, Guo, Simmons et al. 2022). In brief, right-handed non-smokers between 18-45 years of age with a reported BMI between 18.5-24.9 or greater than 30 kg/m^2^ were recruited from the Washington DC metro area. All were free from diabetes, recent weight change (> ± 5 kg in the past 6 months), physical mobility impairments, past or present history of drug abuse, neurological, or psychiatric disorders (including eating disorders such as binge eating) as assessed by an abbreviated Structured Clinical Interview for the Diagnostic and Statistical Manual of Mental Disorders. Furthermore, participants were free from evidence of diseases or medications interfering with study outcomes, allergies to food or local anesthetics, evidence of regular excessive use of caffeinated drinks and alcohol or had strict dietary restrictions. Premenopausal women were studied in the follicular phase for each inpatient visit and were excluded if they were pregnant or breastfeeding. All study procedures were approved by Institutional Review Board participants were compensated for their participation.

#### Fasted cohort

Forty-three adults provided informed consent to participate in an inpatient study of striatal dopamine type 2 receptor binding potential (ClinicalTrials.gov NCT00846040). Study outcomes were reported elsewhere regarding the primary metabolic (Hall et al., 2015) and primary PET outcomes (Guo, Simmons et al. 2014, Guo, Simmons et al. 2022). In brief, right-handed non-smokers between 18-45 years of age with a reported BMI between 18.5-24.9 or greater than 30 kg/m^2^ were recruited from the Washington DC metro area. All were free from diabetes, recent weight change (> ± 5 kg in the past 6 months), physical mobility impairments, past or present history of drug abuse, neurological, or psychiatric disorders (including eating disorders such as binge eating) as assessed by an abbreviated Structured Clinical Interview for the Diagnostic and Statistical Manual of Mental Disorders. Furthermore, participants were free from evidence of diseases or medications interfering with study outcomes, allergies to food or local anesthetics, evidence of regular excessive use of caffeinated drinks and alcohol or had strict dietary restrictions. Premenopausal women were studied in the follicular phase for each inpatient visit and were excluded if they were pregnant or breastfeeding. All study procedures were approved by Institutional Review Board participants were compensated for their participation.

### Diet stabilization

#### Postprandial cohort

For the inpatient portion of their visits, all subjects were confined to the metabolic ward throughout the study without access to outside food. Meals were consumed under observation, and any uneaten food was weighed back to subsequent meals were adjusted to account for uneaten food as needed. Diets were designed using ProNutra software (version 3.4, Viocare, Inc.). The energy-balanced standard diet was based on the resting energy expenditure measured at screening with an activity factor of 1.5 using a standard diet composition of 50% carbohydrate, 35% fat, and 15% protein was consumed for at least one day prior to measuring [^18^F]fallypride D2BP. All participants were provided with a weight maintenance standard diet from the NIH Clinical Center Metabolic Kitchen on an outpatient basis for 2 days prior to admission and at least one full day of admission (≥ 3 full days diet stabilization) before their morning [^18^F]fallypride PET scan two hours after a standardized breakfast reflecting 20% of energy needs.

#### Fasted cohort

Participants were placed on a standard eucaloric diet (50% carbohydrate, 15% protein, 35% fat) with daily energy needs calculated using the Mifflin-St Jeor equation and standard activity factor of 1.5. All meals were prepared in the NIH Clinical Center Nutrition Department Metabolic Kitchen with all foods and beverages weighed on a gram scale (Mettler Toledo Model MS12001L/03). For the run-in phase, participants were provided with 3-5 days of meals and consumed them at home prior to admission to the NIH Clinical Center. Participants were instructed to consume all foods and beverages provided. Any food not consumed was returned and weighed back. For any foods or beverages participants consumed off of the standardized run-in diet, participants were asked to provide a description and amount of what was consumed so that total daily nutrient intake was captured. During the inpatient phase, participants were instructed to consume all foods and beverages provided, and access to outside food was restricted while on the metabolic ward. Any foods or beverages not consumed were weighed back with energy and macronutrients replaced at the next available meal.

### Anthropometrics and body composition

Height was measured in centimeters using a wall stadiometer (Seca 242, Hanover, MD, USA) and weight was measured in kilograms using a digital scale (Scale-Tronix 5702, Carol Steam, IL, USA). All measurements were obtained after an overnight fast while participants were wearing only hospital scrubs.

During the inpatient stay, participants each completed a Dual Energy X-Ray Absorptiometry (DEXA) scan while wearing hospital gown/scrubs to determine body composition (General Electric Lunar iDXA; General Electric; Milwaukee, WI, USA).

### Analytical Measurements and HOMA-IR Calculation

Blood was collected in the overnight fasted state, drawn into chilled EDTA-coated tubes containing preservative (glucose: GLT additive; insulin: SST additive) and kept on ice until centrifuged (1600 *g* for 15 min at 4°C) within 30 min of collection for isolation of plasma. Samples were processed immediately after collection and portions stored for future measurement of biomarkers. Glucose was analyzed using Hexokinase method assayed on Abbott Architect. Insulin was analyzed using electrochemiluminescence Immunoassay on Roche Cobas e601 analyzer. HOMA-IR was calculated as Fasting Insulin (mU/L) x Fasting Glucose (mg/dL)) / 405.

### Resting Energy Expenditure

While inpatient, after a 12 hour overnight fast, participants underwent indirect calorimetry using the ventilated hood technique while supine. Data were collected for 30 minutes, and the first 5 minutes were excluded from analysis. Resting energy expenditure was calculated using the principles of indirect calorimetry using the VO_2_ and VCO_2_ measurements (Weir 1949).

#### Three Factor Eating Questionnaire (TFEQ)

Participants completed a self-assessment questionnaire developed to measure eating behavior traits of dietary restraint, disinhibition and hunger (Stunkard and Messick 1985).

### Magnetic Resonance Imaging

#### Fed cohort

On the afternoon following the morning PET scanning, high resolution anatomical brain MRI was acquired with a HDx General Electric 3 Tesla scanner (TE = 2.7ms, TR 7.24 ms, flip angle 12°, voxel size 0.937*0.937*1.2mm) for each subject.

#### Fasted cohort

During their inpatient stay, high resolution anatomical brain MRI was acquired for each subject. Due to the duration of data collection, extended by the COVID-19 pandemic, T1 weighted structural MRIs were collected on 3T Siemens Verio (n=21; TE = 2.98 ms, TR = 2.3 ms, TI = 900 ms, flip angle 9°, slice thickness = 1.2 mm, voxel size 1*1*1.2mm), and on 3T GE MR-750 Discovery scanner (n=6, TE = 3.04 ms, TR = 7.648 ms, TI = 1060 ms, flip angle 8°, slice thickness = 1.0 mm, voxel size 1*1*1mm; n=32, TE= 3.46 ms, TR = 8.156 ms, TI = 900 ms, flip angle 7°, slice thickness = 1.0 mm, voxel size 1*1*1 mm) for each subject. Quality of individual subject data were checked by study team [VLD & JG].

The anatomical images were parcellated with FreeSurfer software to generate ROI binary mask volumes in each subject in the putamen, caudate, accumbens, pallidum, and the cerebellum (reference region) (http://surfer.nmr.mgh.harvard.edu). All individual ROI masks were visually checked.

#### Positron Emission Tomography

The Fed cohort completed [^18^F]fallypride scans starting two hours after a standardized breakfast (0800 hours) comprising 20% of energy needs as estimated using indirect calorimetry, while the Fasted cohort completed morning [^18^F]fallypride scans in the confirmed overnight fasted (15 hours) state.

All PET scanning was performed using Siemens HRRT (Siemens Healthcare, Malvern, PA) immediately after a bolus of 5 mCi of [^18^F]fallypride infused intravenously using a Harvard® pump. The radiochemical purity of the radiotracer was > 99%. PET emission data were collected over 3.5 hours in three blocks separated by two 10-minute breaks. Thirty-three volumes were acquired at times 0, 0.25, 0.5, 0.75, 1, 1.25, 1.5, 1.75, 2, 2.5, 3, 3.5, 4, 4.5, 5, 6, 7, 8, 9, 10, 12.5, 15, 20, 25, 30, 40, 50, 60, 90, 110, 130, 170, 200 min. During each scan block, the room was quiet and dimly lit and each subject was instructed to keep their head as still as possible, relax, and try to avoid falling asleep. The image reconstruction process were corrected for head motion which was tracked throughout each scan.

Each scan consisted of 207 slices (slice separation = 1.22 mm). The fields of view were 31.2 cm and 25.2 cm for transverse and axial slices, respectively. A transmission scan was obtained with a 137Cs rotating pin source before radiotracer injection and before the emission scan to correct for attenuation. The PET images were aligned within each scan block with 6-parameter rigid registration using 7th order polynomial interpolation and each block was aligned to the volume taken at 20 min of the first block. The final alignments were visually checked, with translations varying by <5 mm and the rotations by <5 degrees.

<u>For region of interest analyses,</u> individual participants’ anatomical MRI images were co-registered to the aligned PET images by minimizing a mutual information cost function. The final coregistration was visually checked. Time-activity curves within the Freesurfer-generated ROIs were extracted and kinetic parameters were fit to a two-compartment model to determine regional D2BP (i.e., BP*_ND_* (Lammertsma and Hume 1996, Innis, Cunningham et al. 2007). Two-sided t-tests were used to compare the mean dopamine D2 receptor non-displaceable binding potential (BP*_ND_*; referred to here as D2BP) within each ROI between the fasted and fed states.

For <u>voxelwise analyses, e</u>ach individual’s co-registered anatomical MRI was linearly transformed into the Talairach space, and the transformation matrix was applied to the PET images which were then smoothed with a 5-mm full-width, half-max Gaussian kernel. Data were exported from Talairach space to MATLAB where time-activity curves for tracer concentration in each voxel were fit to the same two compartment model (as above) using the cerebellum as a reference tissue to determine D2BP at each voxel and exported back to Talairach space for group level spatial analyses. The individual subject’s D2BP maps were included in a regression analysis in AFNI’s 3dttest++ to identify clusters where D2BP across subjects were correlated with BMI. As high D2BP occurs mainly in striatum, small volume corrections were implemented within striatum where D2BP is > 1.5. A two-sided voxel-wise threshold of P < 0.1 was used, and cluster size threshold to achieve correction for multiple comparisons at P < 0.05 was implemented via Monte Carlo simulations in AFNI’s 3dClustSim. When comparing voxelwise D2BP, the analyses were expanded to encompass the whole brain. Voxel-wise threshold was set at *p-*uncorrected<0.0001 and cluster size threshold at 30 voxels, and FDR corrected<0.001.

## Declaration of generative AI and AI-assisted technologies in the manuscript preparation process

During the preparation of this work, the author(s) used **GPT-5.6** for light editing. The author(s) reviewed and edited the output as needed and take full responsibility for the content of the published article.

## SUPPLEMENTARY INFORMATION

**Supplementary Figure 1.**
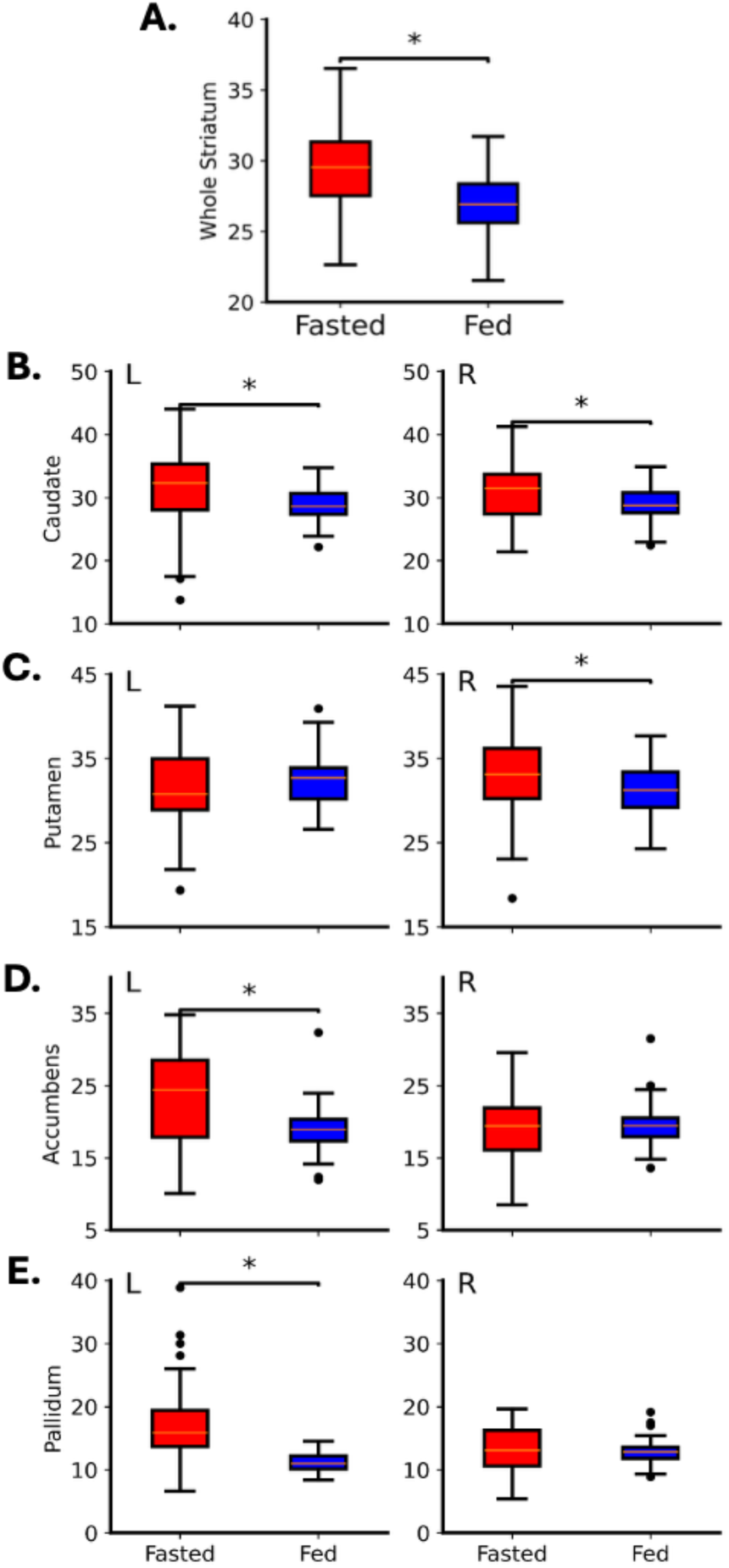
Comparisons of striatal dopamine D2R binding potential in ROIs between Fed vs Fasted Cohorts adjusted for age, sex, and HOMA-IR. Dopamine D2R binding potential measured by [18F]fallypride is significantly reduced in individuals in the 2-hour post-ingestive state (Fed, n=43; blue bars) compared to individuals in the overnight fasted state (Fasted n=57; red bars) The differences in binding potential (mean±SEM) were (A) whole striatum, −2.93 ± 0.68 (p < 0.0001), (B) caudate (left, −2.5 ± 1.06 (p = 0.02); right, −2.01 ± 0.9 (p = 0.03)), (C) putamen (left, 1 ± 0.93 (p = 0.29); right, −1.93 ± 0.96 (p = 0.047)), (D) accumbens (left, −4.36 ± 1.21 (p = 0.0006); right, 0.77 ± 0.96 (p = 0.42)) and (E) pallidum (left, −5.86 ± 1.03 (p < 0.001); right, −0.21 ± 0.66 (p = 0.75)).

**Supplementary figure 2.**
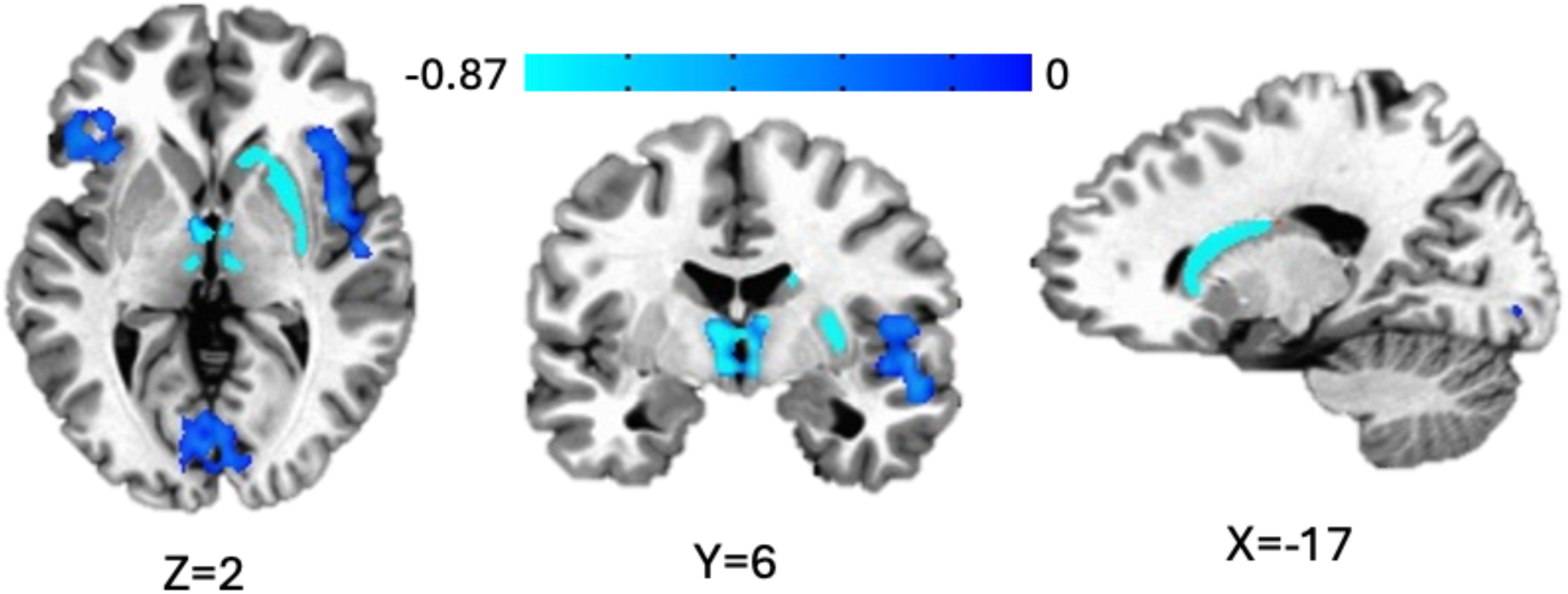
Voxel-wise comparisons of striatal dopamine D2R binding in fed vs. fasting state cohorts after adjusting for age, sex, and insulin sensitivity (HOMA-IR). Voxel-wise analyses identified clusters across the whole brain where dopamine D2R binding potentials significantly decreased in fed state compared to fasting state Clusters are defined by faces-touching neighbor clustering, bi-sided uncorrected p < 0.0001, cluster size > 30 voxels, and multiple-comparison corrected p < 0.05. Voxels are displayed after cubic convolution resampling. The color map represents the reduced BP’s in individual voxels.

**Supplementary figure 3.**
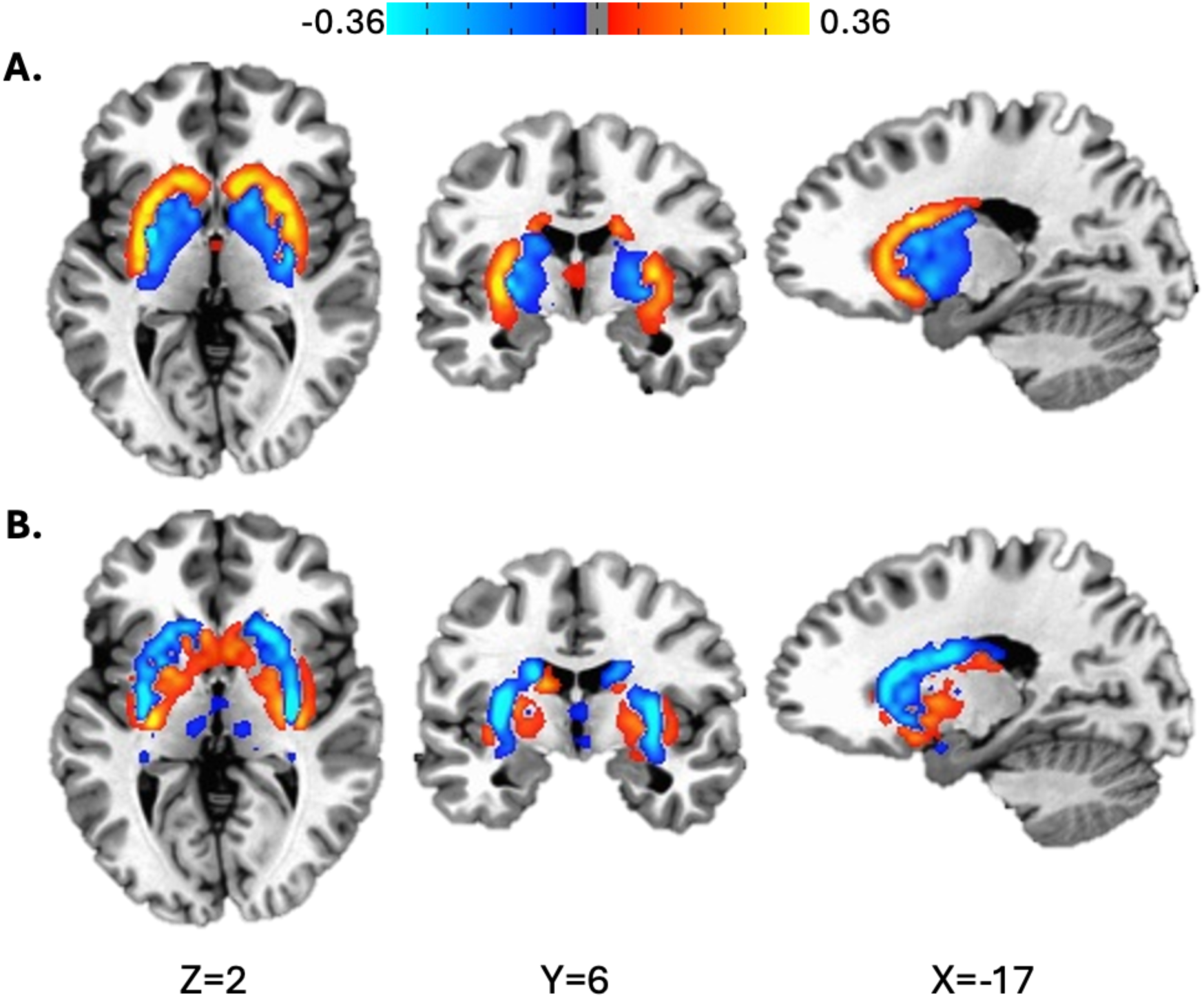
Unthresholded voxel-wise correlation patterns between dopamine D2R binding and BMI in **(A)** Fed Cohort and **(B)** Fasted Cohort. Regions where D2BP increases with BMI (suggestive of reduced synaptic dopamine) indicated in red/orange, while regions where D2BP decreases with BMI (suggestive of increased synaptic dopamine) indicated in blues. Clusters are defined by faces-touching neighbor clustering, bi-sided uncorrected p < 0.0001, cluster size > 30 voxels. Clusters uncorrected for multiple comparisons for visualization purposes. Voxels are displayed after cubic convolution resampling.

**Supplementary figure 4.**
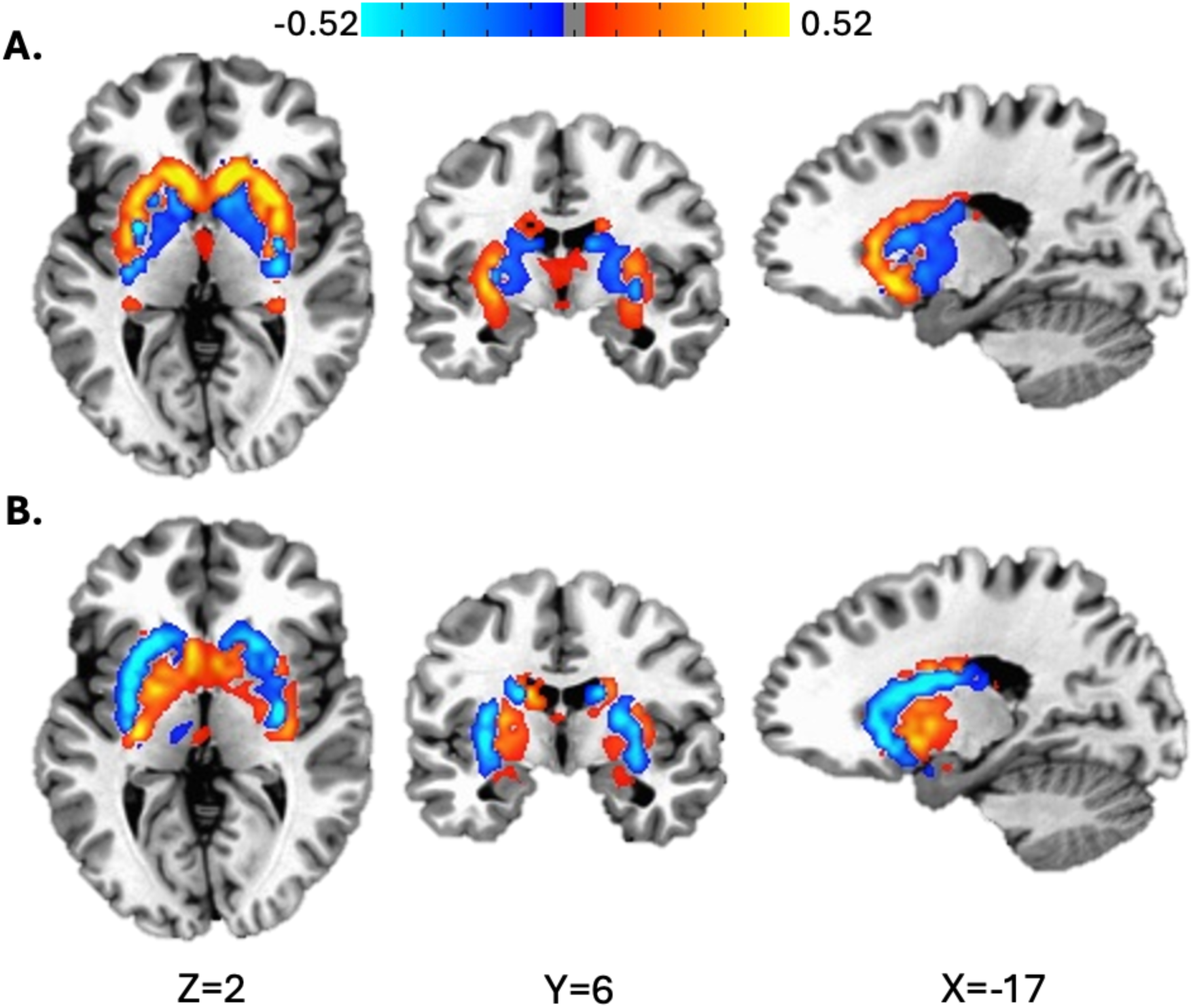
Unthresholded voxel-wise correlation patterns between dopamine D2R binding and BMI *adjusted for age, sex, and HOMA-IR* in **(A)** Fed Cohort and **(B)** Fasted Cohort. Regions where D2BP increases with BMI (suggestive of reduced synaptic dopamine) indicated in red/orange, while regions where D2BP decreases with BMI (suggestive of increased synaptic dopamine) indicated in blues. Clusters are defined by faces-touching neighbor clustering, bi-sided uncorrected p < 0.0001, cluster size > 30 voxels. Clusters uncorrected for multiple comparisons for visualization purposes. Voxels are displayed after cubic convolution resampling.

**Supplementary figure 5A-E.**
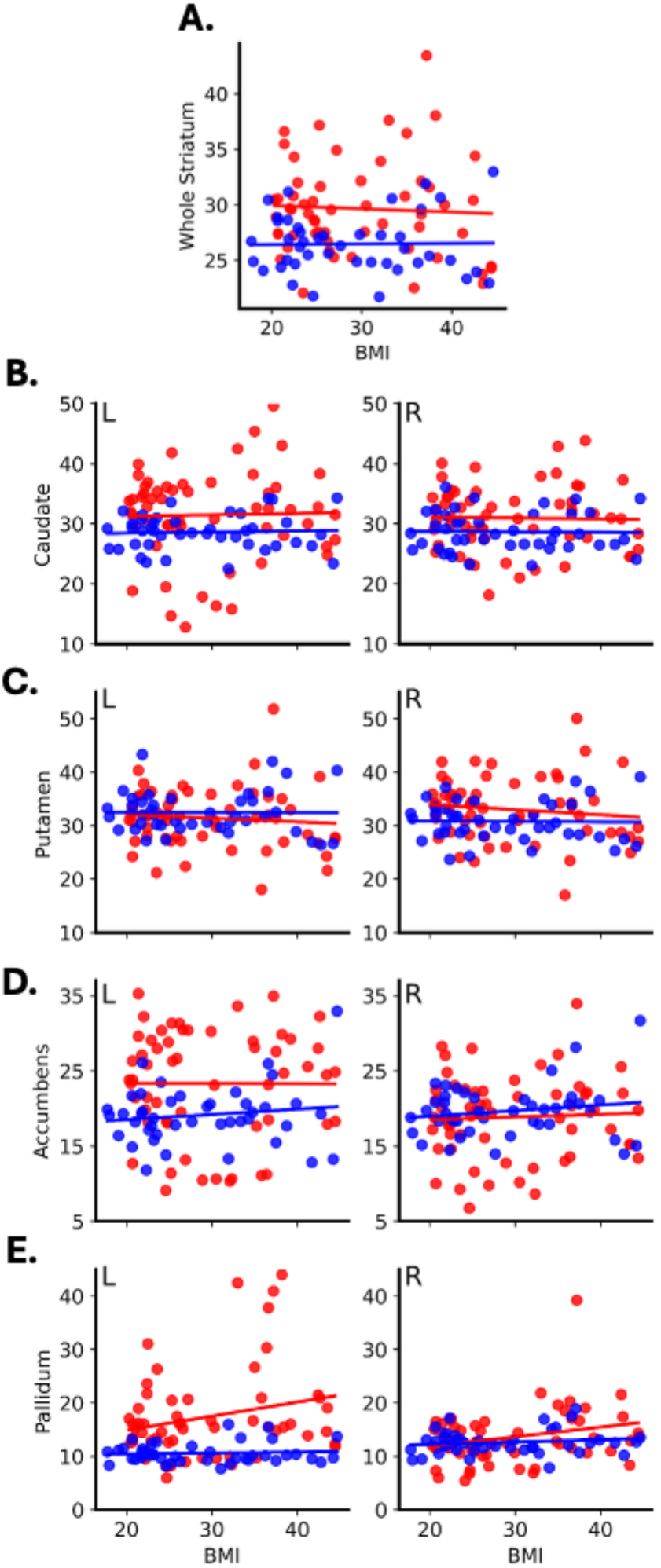
Similar relationship between BMI and D2BP in ROIs between fed and fasting states. Fasted cohort (red); fed cohort (blue). Region of interest analysis comparing slopes of the relationship between D2BP as measured by [18F]fallypride and adiposity as measured by body mass index (BMI; kg/m2). Regression coefficients at the level of (A) the whole striatum were β_fed_ vs. β_fasting_ were 0.006 ± 0.074 vs. −0.029 ± 0.066 (p = 0.72). Regression coefficients across striatal subregions β_fed_ vs. β_fasting_ were as follows: (B) left caudate: 0.018 ± 0.121vs 0.025 ± 0.108 (p = 0.97); right caudate: −0.006 ± 0.094 vs. −0.015 ± 0.083 (p = 0.94); (C) left putamen: −0.002 ± 0.1 vs. −0.068 ± 0.089 (p = 0.62); right putamen: −0.008 ± 0.102 vs. −0.095 ± 0.091 (p = 0.52); (D) left accumbens: 0.07 ± 0.121 vs. −0.002 ± 0.107 (p = 0.66); right accumbens: 0.074 ± 0.095 vs. 0.036 ± 0.085 (p = 0.77); (E) left pallidum: 0.019 ± 0.127 vs. 0.263 ± 0.113 (p = 0.15); right pallidum: 0.042 ± 0.082 vs. 0.181 ± 0.073 (p = 0.21).

**Supplementary figure 6A-E.**
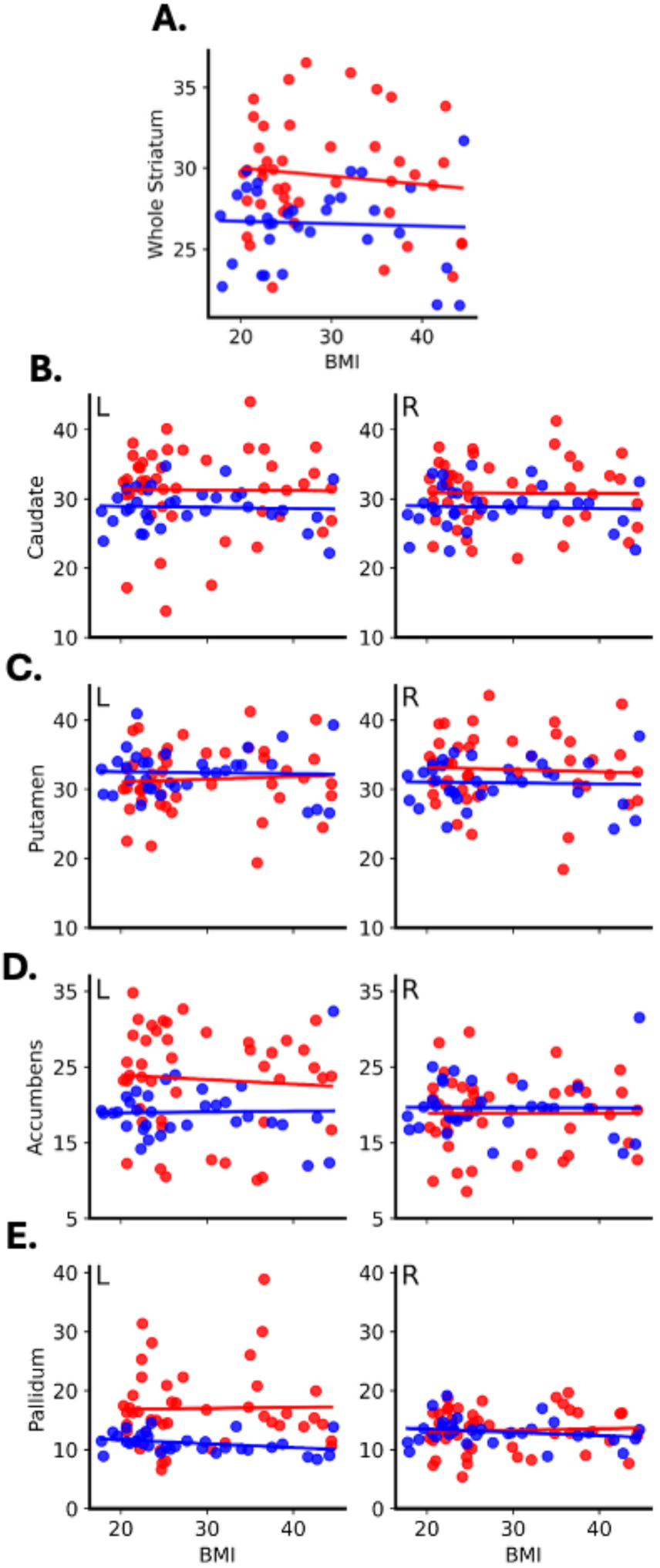
Similar relationship between BMI and D2BP in ROIs between cohorts in fed and fasting states, adjusted for age, sex and insulin sensitivity. Fasted cohort (red); fed cohort (blue). Region of interest analysis comparing slopes of the relationship between D2BP as measured by [18F]fallypride and adiposity as measured by body mass index (kg/m2), adjusted for age, sex and insulin sensitivity (HOMA-IR). Regression coefficients at the level of (A) the whole striatum were β_fed_ vs. β_fasting_ were −0.001 ± 0.073 vs −0.029 ± 0.077 (p = 0.77). Regression coefficients β_fed_ vs. β_fasting_ across striatal subregions were as follows: (A) left caudate: 0.005 ± 0.121 vs. 0.029 ± 0.128 (p = 0.88); right caudate: −0.001 ± 0.099 vs. 0.027 ± 0.104 (p = 0.83); (B) left putamen −0.015 ± 0.102 vs. 0.04 ± 0.108 (p = 0.68); right putamen: −0.004 ± 0.106 vs. −0.019 ± 0.112 (p = 0.92); (C) left accumbens: 0.039 ± 0.136 vs. −0.013 ± 0.143 (p = 0.77); right accumbens: −0.011 ± 0.105 vs. −0.015 ± 0.11 (p = 0.98); (D) left pallidum: −0.008 ± 0.12 vs. 0.123 ± 0.126 (p = 0.41); and right pallidum: −0.052 ± 0.074 vs. 0.034 ± 0.078 (p = 0.38).

**Supplementary Figure 7.**
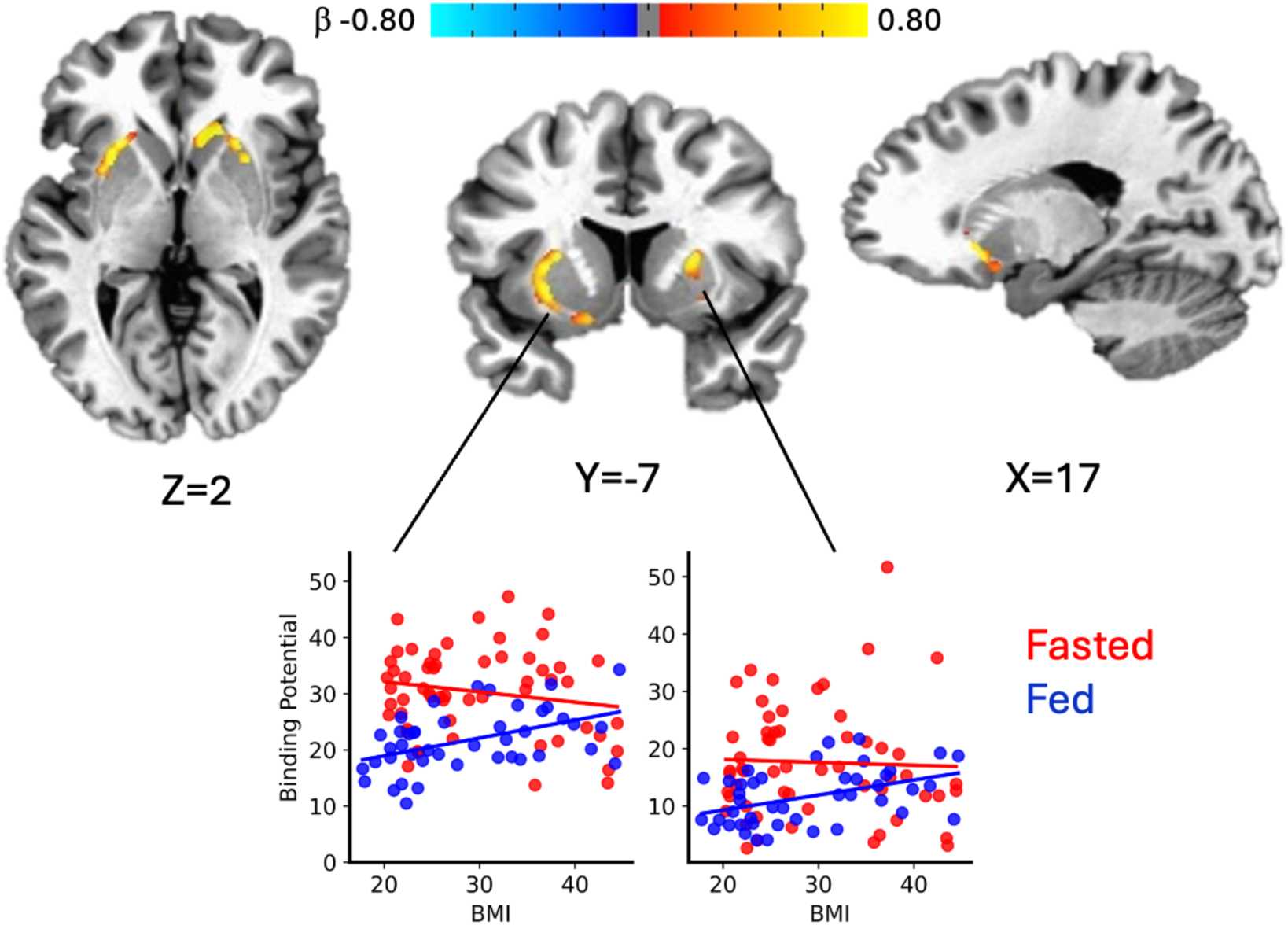
Voxelwise comparison of correlation between dopamine D2 binding potential and adiposity in Fed and Fasted cohorts, adjusted for age, sex, and insulin sensitivity. Voxel-wise analyses within striatal mask identified clusters where the regression β coefficients of the D2BP-BMI relationship are significantly different in fed vs fasted cohorts in the left dorsolateral putamen (left plot) and in the right pallidum (right plot). Clusters are defined by faces-touching neighbor clustering, bi-sided uncorrected p < 0.1, cluster size > 30 voxels, and multiple-comparison corrected p < 0.05. Voxels are displayed after cubic convolution resampling. The color maps represent the difference of the regression β-coefficients between the fed and fasting states, displayed in cluster-specific scatter plots for visualization purposes.

---

**Supplementary Table 1.** Locations of clusters from voxelwise analyses after adjusting for age sex and insulin sensitivity (HOMA-IR). PET resolution 3.5mm^3^. Imaging analyses conducted in Analysis of Functional Neuroimaging (AFNI) (clusters defined by voxels with faces touching). Coordinates in Talairach Space.

|  | Location of Peak |  |  | Voxels | Size<br>(mm3) | t-stat | alpha |
| --- | --- | --- | --- | --- | --- | --- | --- |
|  | x | y | z |  |  |  |  |
| **Comparisons of BPs adjusted for age sex and HOMA** |  |  |  |  |  |  |  |
| Right insula | -54.2 | 18.5 | -4 | 170 | 7289 | -4.45 | <0.001 |
| Right Inferior frontal gyrus<br>(pars Orbitalis) |  |  |  |  |  |  |  |
| Right superior temporal<br>gyrus |  |  |  |  |  |  |  |
| Left calcarine gyrus | 1.8 | 78 | 3 | 117 | 5016 | -6.71 | <0.001 |
| Right calcarine gyrus |  |  |  |  |  |  |  |
| Left lingual gyrus |  |  |  |  |  |  |  |
| Left inferior frontal gyrus<br>(pars Triangularis) | 47.2 | -23.5 | 3 | 84 | 3602 | -6.74 | <0.001 |
| Left inferior frontal gyrus<br>(pars Orbitalis) |  |  |  |  |  |  |  |
| Left insula |  |  |  |  |  |  |  |
| Left thalamus | 1.8 | 8 | 6.5 | 51 | 2187 | -7.85 | <0.001 |
| Right thalamus |  |  |  |  |  |  |  |
| Left ventral diencephalon |  |  |  |  |  |  |  |
| Right putamen | -29.8 | 8 | 3 | 36 | 1544 | -5.99 | <0.001 |
| Right pallidum |  |  |  |  |  |  |  |
| Right caudate | -15.8 | -13.0 | 17 | 32 | 1372 | -6.37 | <0.001 |
| Right putamen |  |  |  |  |  |  |  |
| **Comparisons of BMI slopes adjusted for age sex and HOMA** |  |  |  |  |  |  |  |
| Right caudate | -15.8 | -20 | 6.5 | 57 | 2444 | 2.70 | <0.005 |
| Right putamen |  |  |  |  |  |  |  |
| Right olfactory cortex |  |  |  |  |  |  |  |
| Left putamen | 19.2 | -16.5 | -0.5 | 48 | 2058 | 2.58 | <0.005 |
| Left insula |  |  |  |  |  |  |  |
| Left olfactory cortex |  |  |  |  |  |  |  |

|  | Location of Peak |  |  | Voxels | Size | t-stat | alpha |
| --- | --- | --- | --- | --- | --- | --- | --- |
|  | x | y | z |  | (mm3) |  |  |
| **Difference Between Cohorts, adjusted for age, sex and HOMA-IR (D2BP Fed – D2BP Fasted)<sup>#</sup>** |  |  |  |  |  |  |  |
| Insula, right | -54.2 | 18.5 | -4 | 170 | 7289 | -4.45 | <0.001 |
| Inferior frontal gyrus (pars Orbitalis), right |  |  |  |  |  |  |  |
| Superior temporal gyrus, right |  |  |  |  |  |  |  |
| Calcarine gyrus, left | 1.8 | 78 | 3 | 117 | 5016 | -6.71 | <0.001 |
| Calcarine gyrus, right |  |  |  |  |  |  |  |
| Lingual gyrus, left |  |  |  |  |  |  |  |
| Inferior frontal gyrus (pars Triangularis), left | 47.2 | -23.5 | 3 | 84 | 3602 | -6.74 | <0.001 |
| Inferior frontal gyrus (pars Orbitalis), left |  |  |  |  |  |  |  |
| Insula, left |  |  |  |  |  |  |  |
| Thalamus, left | 1.8 | 8 | 6.5 | 51 | 2187 | -7.85 | <0.001 |
| Thalamus, right |  |  |  |  |  |  |  |
| Ventral diencephalon, left |  |  |  |  |  |  |  |
| Putamen, right | -29.8 | 8 | 3 | 36 | 1544 | -5.99 | <0.001 |
| Pallidum, right |  |  |  |  |  |  |  |
| Caudate, right | -15.8 | -13.0 | 17 | 32 | 1372 | -6.37 | <0.001 |
| Putamen, right |  |  |  |  |  |  |  |
| **Comparisons of BMI slopes adjusted for age sex and HOMA-IR ($\beta$ Fed – $\beta$ Fasted)<sup>\*</sup>** |  |  |  |  |  |  |  |
| Caudate, right | -15.8 | -20 | 6.5 | 57 | 2444 | 2.70 | <0.005 |
| Putamen, right |  |  |  |  |  |  |  |
| Olfactory cortex, right |  |  |  |  |  |  |  |
| Putamen, left | 19.2 | -16.5 | -0.5 | 48 | 2058 | 2.58 | <0.005 |
| Insula, left |  |  |  |  |  |  |  |
| Olfactory cortex, left |  |  |  |  |  |  |  |

Maximum of top three descriptive labels reflecting largest proportion of cluster provided using the atlas accounting for greatest percent of anatomical localization of cluster.
<sup>#</sup> Clusters are defined by faces-touching neighbor clustering, bi-sided uncorrected $p < 0.0001$ , cluster size $> 30$ voxels, and multiple-comparison corrected $p < 0.05$ .
<sup>\*</sup> Clusters are defined by faces-touching neighbor clustering, bi-sided uncorrected $p < 0.1$ , cluster size $> 30$ voxels, and multiple-comparison corrected $p < 0.05$ .

## REFERENCES

Abizaid, A., Z.-W. Liu, Z. B. Andrews, M. Shanabrough, E. Borok, J. D. Elsworth, R. H. Roth, M. W. Sleeman, M. R. Picciotto and M. H. Tschöp (2006). “Ghrelin modulates the activity and synaptic input organization of midbrain dopamine neurons while promoting appetite.” The Journal of clinical investigation 116(12): 3229–3239.

Avery, J. A., K. L. Kerr, J. E. Ingeholm, K. Burrows, J. Bodurka and W. K. Simmons (2015). “A common gustatory and interoceptive representation in the human mid-insula.” Human Brain Mapping 36(8): 2996–3006.

Banks, W., S. Farr, T. Salameh, M. Niehoff, E. Rhea, J. Morley, A. Hanson, K. Hansen and S. Craft (2018). “Triglycerides cross the blood–brain barrier and induce central leptin and insulin receptor resistance.” International journal of obesity 42(3): 391–397.

Berland, C., E. Montalban, E. Perrin, M. Di Miceli, Y. Nakamura, M. Martinat, M. Sullivan, X. S. Davis, M. A. Shenasa, C. Martin, S. Tolu, F. Marti, S. Caille, J. Castel, S. Perez, C. G. Salinas, C. Morel, J. Hecksher-Sørensen, M. Cador, X. Fioramonti, M. H. Tschöp, S. Layé, L. Venance, P. Faure, T. S. Hnasko, D. M. Small, G. Gangarossa and S. H. Luquet (2020). “Circulating Triglycerides Gate Dopamine-Associated Behaviors through DRD2-Expressing Neurons.” Cell Metabolism 31(4): 773–790.e711.

Berry, S. E., A. M. Valdes, D. A. Drew, F. Asnicar, M. Mazidi, J. Wolf, J. Capdevila, G. Hadjigeorgiou, R. Davies, H. Al Khatib, C. Bonnett, S. Ganesh, E. Bakker, D. Hart, M. Mangino, J. Merino, I. Linenberg, P. Wyatt, J. M. Ordovas, C. D. Gardner, L. M. Delahanty, A. T. Chan, N. Segata, P. W. Franks and T. D. Spector (2020). “Human postprandial responses to food and potential for precision nutrition.” Nat Med 26(6): 964–973.

Bozzetto, L., G. Della Pepa, C. Vetrani and A. A. Rivellese (2020). “Dietary Impact on Postprandial Lipemia.” Frontiers in Endocrinology 11: 337–337.

Cansell, C., J. Castel, R. Denis, C. Rouch, A. S. Delbes, S. Martinez, D. Mestivier, B. Finan, J. G. Maldonado-Aviles, M. Rijnsburger, M. H. Tschöp, R. J. Dileone, R. H. Eckel, L. Fleur, C. Magnan, T. S. Hnasko and S. Luquet (2014). “Dietary triglycerides act on mesolimbic structures to regulate the rewarding and motivational aspects of feeding.” Molecular Psychiatry 19: 1095–1105.

Carnell, S., K. E. Steele, G. Thapaliya, H. Kuwubara, A. Aghababian, A. Papantoni, A. Nandi, J. R. Brašić, T. H. Moran and D. F. Wong (2023). “Milkshake Acutely Stimulates Dopamine Release in Ventral and Dorsal Striatum in Healthy-Weight Individuals and Patients with Severe Obesity Undergoing Bariatric Surgery: A Pilot Study.” Nutrients 15(12): 2671.

Carr, K. D. (2007). “Chronic food restriction: Enhancing effects on drug reward and striatal cell signaling.” Physiology & Behavior 91(5): 459–472.

Ceccarini, J., E. Vrieze, M. Koole, T. Muylle, G. Bormans, S. Claes and K. Van Laere (2012). “Optimized in vivo detection of dopamine release using 18F-fallypride PET.” J Nucl Med 53(10): 1565–1572.

Cone, J. J., J. E. McCutcheon and M. F. Roitman (2014). “Ghrelin acts as an interface between physiological state and phasic dopamine signaling.” J Neurosci 34(14): 4905–4913.

Cropley, V. L., R. B. Innis, P. J. Nathan, A. K. Brown, J. L. Sangare, A. Lerner, Y. H. Ryu, K. E. Sprague, V. W. Pike and M. Fujita (2008). “Small effect of dopamine release and no effect of dopamine depletion on [18F]fallypride binding in healthy humans.” Synapse 62(6): 399–408.

Dang, L. C., G. R. Samanez-Larkin, J. J. Castrellon, S. F. Perkins, R. L. Cowan and D. H. Zald (2016). “Associations between dopamine D2 receptor availability and BMI depend on age.” NeuroImage 138: 176–183.

Darcey, V. L., J. Guo, M. Chi, S. T. Chung, A. B. Courville, I. Gallagher, P. Herscovitch, R. Howard, M. La Noire, L. Milley, A. Schick, M. Stagliano, S. Turner, N. Urbanski, S. Yang, E. Yim, N. Zhai, M. S. Zhou and K. D. Hall (2025). “Striatal dopamine tone is positively associated with adiposity in humans as determined by PET using dual dopamine type-2 receptor antagonist tracers.” Molecular Psychiatry 30(8): 3708–3717.

Darcey, V. L., J. Guo, M. Chi, S. T. Chung, A. B. Courville, I. Gallagher, P. Herscovitch, R. Howard, M. LaNoire, L. Milley, A. Schick, M. Stagliano, S. Turner, N. Urbanski, S. Yang, E. Yim, N. Zhai, M. S. Zhou and K. D. Hall (2023). “Striatal dopamine tone is positively associated with body mass index in humans as determined by PET using dual dopamine type-2 receptor antagonist tracers.” <u>medRxiv</u>: 2023.2009.2027.23296169.

Darcey, V. L., J. Guo, M. Chi, S. T. Chung, A. B. Courville, I. Gallagher, P. Herscovitch, P. V. Joseph, R. Howard, M. La Noire, L. Milley, A. Schick, M. Stagliano, S. Turner, N. Urbanski, S. Yang, N. Zhai, M. S. Zhou and K. D. Hall (2025). “Brain dopamine responses to ultra-processed milkshakes are highly variable and not significantly related to adiposity in humans.” Cell Metab 37(3): 616–628.e615.

de Araujo, I. E., J. G. Ferreira, L. A. Tellez, X. Ren and C. W. Yeckel (2012). “The gut-brain dopamine axis: a regulatory system for caloric intake.” Physiol Behav 106(3): 394–399.

de Araujo, I. E., M. Schatzker and D. M. Small (2020). “Rethinking Food Reward.” Annu Rev Psychol 71: 139–164.

DiFeliceantonio, A. G. and D. M. Small (2019). “Dopamine and diet-induced obesity.” Nature Neuroscience 22(1): 1–2.

Eisenstein, S. A., K. J. Black, A. Samara, J. M. Koller, J. P. Dunn, T. Hershey, S. Klein and G. I. Smith (2020). “Striatal Dopamine Responses to Feeding are Altered in People with Obesity.” Obesity (Silver Spring, Md.) 28(4): 765–765.

Farooqi, I. S., E. Bullmore, J. Keogh, J. Gillard, S. O’Rahilly and P. C. Fletcher (2007). “Leptin regulates striatal regions and human eating behavior.” Science 317(5843): 1355.

Ferreira, J. G., L. A. Tellez, X. Ren, C. W. Yeckel and I. E. de Araujo (2012). “Regulation of fat intake in the absence of flavour signalling.” The Journal of physiology 590(4): 953–972.

Ford, C. P. (2014). “The role of D2-autoreceptors in regulating dopamine neuron activity and transmission.” Neuroscience 282: 13–22.

Gautier, J. F., K. Chen, A. D. Salbe, D. Bandy, R. E. Pratley, M. Heiman, E. Ravussin, E. M. Reiman and P. A. Tataranni (2000). “Differential brain responses to satiation in obese and lean men.” Diabetes 49(5): 838–846.

Grace, A. A. (2000). “The tonic/phasic model of dopamine system regulation and its implications for understanding alcohol and psychostimulant craving.” Addiction 95 **Suppl 2**: S119–128.

Grove, J. C. R., A. M. Hakimi, Q. Li, J. Zhang, H. Backes, B. Kuzmanovic, J. Choi, V. Ubadiah, L. Qiu, Z. Liu, D. M. Small, M. Tittgemeyer and Z. A. Knight (2025). “A lateralized pathway for associating nutrients with flavors.” bioRxiv.

Guitart-Masip, M., Q. J. Huys, L. Fuentemilla, P. Dayan, E. Duzel and R. J. Dolan (2012). “Go and no-go learning in reward and punishment: interactions between affect and effect.” Neuroimage 62(1): 154–166.

Guo, J., W. K. Simmons, P. Herscovitch, A. Martin and K. D. Hall (2014). “Striatal dopamine D2-like receptor correlation patterns with human obesity and opportunistic eating behavior.” Molecular Psychiatry 19(10): 1078–1084.

Guo, J., W. K. Simmons, P. Herscovitch, A. Martin and K. D. Hall (2022). “Correction to: Striatal dopamine D2-like receptor correlation patterns with human obesity and opportunistic eating behavior.” Mol Psychiatry 27(10): 4369.

Hall, K. D. and V. L. Darcey (2025). “The peril of preconceived narratives.” Cell Metab 37(8): 1624–1625.

Han, W., Luis A. Tellez, J. Niu, S. Medina, Tatiana L. Ferreira, X. Zhang, J. Su, J. Tong, Gary J. Schwartz, A. van den Pol and Ivan E. de Araujo (2016). “Striatal Dopamine Links Gastrointestinal Rerouting to Altered Sweet Appetite.” Cell Metabolism 23(1): 103–112.

Heni, M. (2024). “The insulin resistant brain: impact on whole-body metabolism and body fat distribution.” Diabetologia 67(7): 1181–1191.

Holroyd, C. B., S. Nieuwenhuis, N. Yeung, L. Nystrom, R. B. Mars, M. G. H. Coles and J. D. Cohen (2004). “Dorsal anterior cingulate cortex shows fMRI response to internal and external error signals.” Nature Neuroscience 7(5): 497–498.

Innis, R. B., V. J. Cunningham, J. Delforge, M. Fujita, A. Gjedde, R. N. Gunn, J. Holden, S. Houle, S.-C. Huang, M. Ichise, H. Iida, H. Ito, Y. Kimura, R. A. Koeppe, G. M. Knudsen, J. Knuuti, A. A. Lammertsma, M. Laruelle, J. Logan, R. P. Maguire, M. A. Mintun, E. D. Morris, R. Parsey, J. C. Price, M. Slifstein, V. Sossi, T. Suhara, J. R. Votaw, D. F. Wong and R. E. Carson (2007). “Consensus Nomenclature for in vivo Imaging of Reversibly Binding Radioligands.” Journal of Cerebral Blood Flow & Metabolism 27(9): 1533–1539.

Kessler, R. M., W. O. Whetsell, M. Sib Ansari, J. R. Votaw, T. de Paulis, J. A. Clanton, D. E. Schmidt, N. Scott Mason and R. G. Manning (1993). “Identification of extrastriatal dopamine D2 receptors in post mortem human brain with [125I]epidepride.” Brain Research 609(1): 237–243.

Kumar, M., Y. Wu, J. Knapp, C. L. Pontius, D. Park, R. E. Witte, R. McAllister, K. Gupta, K. N. Rajagopalan, P. De Camilli and T. A. Ryan (2025). “Triglycerides are an important fuel reserve for synapse function in the brain.” Nature Metabolism 7(7): 1392–1403.

Lammertsma, A. A. and S. P. Hume (1996). “Simplified reference tissue model for PET receptor studies.” Neuroimage 4(3 Pt 1): 153–158.

Logan, J., J. S. Fowler, N. D. Volkow, G. J. Wang, Y. S. Ding and D. L. Alexoff (1996). “Distribution volume ratios without blood sampling from graphical analysis of PET data.” J Cereb Blood Flow Metab 16(5): 834–840.

Mamo, J. C., G. F. Watts, P. H. Barrett, D. Smith, A. P. James and S. Pal (2001). “Postprandial dyslipidemia in men with visceral obesity: an effect of reduced LDL receptor expression?” Am J Physiol Endocrinol Metab 281(3): E626–632.

McCutcheon, R. A., M. M. Nour, T. Dahoun, S. Jauhar, F. Pepper, P. Expert, M. Veronese, R. A. Adams, F. Turkheimer, M. A. Mehta and O. D. Howes (2019). “Mesolimbic Dopamine Function Is Related to Salience Network Connectivity: An Integrative Positron Emission Tomography and Magnetic Resonance Study.” Biological Psychiatry 85(5): 368–378.

McDougle, M., A. de Araujo, A. Singh, M. Yang, I. Braga, V. Paille, R. Mendez-Hernandez, M. Vergara, L. N. Woodie, A. Gour, A. Sharma, N. Urs, B. Warren and G. de Lartigue (2024). “Separate gut-brain circuits for fat and sugar reinforcement combine to promote overeating.” Cell Metab 36(2): 393–407.e397.

Miederer, I., H.-G. Buchholz, L. Rademacher, C. Eckart, D. Kraft, M. Piel, C. J. Fiebach and M. Schreckenberger (2025). “Dopaminergic Mechanisms of Cognitive Flexibility: An [18F]Fallypride PET Study.” Journal of Nuclear Medicine 66(3): 405–409.

Mukherjee, J., B. T. Christian, K. A. Dunigan, B. Shi, T. K. Narayanan, M. Satter and J. Mantil (2002). “Brain imaging of 18F-fallypride in normal volunteers: blood analysis, distribution, test-retest studies, and preliminary assessment of sensitivity to aging effects on dopamine D-2/D-3 receptors.” Synapse 46(3): 170–188.

O’Doherty, J. P., R. Deichmann, H. D. Critchley and R. J. Dolan (2002). “Neural Responses during Anticipation of a Primary Taste Reward.” Neuron 33(5): 815–826.

Obradovic, M., E. Sudar-Milovanovic, S. Soskic, M. Essack, S. Arya, A. J. Stewart, T. Gojobori and E. R. Isenovic (2021). “Leptin and Obesity: Role and Clinical Implication.” Front Endocrinol (Lausanne) 12: 585887.

Olsson, H. and L. Farde (2001). “Potentials and pitfalls using high affinity radioligands in PET and SPET determinations on regional drug induced D2 receptor occupancy—a simulation study based on experimental data.” Neuroimage 14(4): 936–945.

Patel, J. C., K. D. Carr and M. E. Rice (2023). “Actions and Consequences of Insulin in the Striatum.” Biomolecules 13(3).

Riccardi, P., R. Li, M. S. Ansari, D. Zald, S. Park, B. Dawant, S. Anderson, M. Doop, N. Woodward, E. Schoenberg, D. Schmidt, R. Baldwin and R. Kessler (2006). “Amphetamine-induced displacement of [18F] fallypride in striatum and extrastriatal regions in humans.” Neuropsychopharmacology: official publication of the American College of Neuropsychopharmacology 31(5): 1016–1026.

Riccardi, P., D. Zald, R. Li, S. Park, M. S. Ansari, B. Dawant, S. Anderson, N. Woodward, D. Schmidt, R. Baldwin and R. Kessler (2006). “Sex differences in amphetamine-induced displacement of [ 18F]fallypride in striatal and extrastriatal regions: A PET study.” American Journal of Psychiatry 163(9): 1639–1641.

Rolls, E. T. and F. Grabenhorst (2008). “The orbitofrontal cortex and beyond: From affect to decision-making.” Progress in Neurobiology 86: 216–244.

Rominger, A., E. Wagner, E. Mille, G. Boning, M. Esmaeilzadeh, B. Wangler, F.-J. Gildehaus, S. Nowak, A. Bruche, K. Tatsch, P. Bartenstein and P. Cumming (2010). “Endogenous competition against binding of [(18)F]DMFP and [(18)F]fallypride to dopamine D(2/3) receptors in brain of living mouse.” Synapse (New York, N.Y.) 64(4): 313–322.

Sander, C. Y., J. M. Hooker, C. Catana, M. D. Normandin, N. M. Alpert, G. M. Knudsen, W. Vanduffel, B. R. Rosen and J. B. Mandeville (2013). “Neurovascular coupling to D2/D3 dopamine receptor occupancy using simultaneous PET/functional MRI.” Proceedings of the National Academy of Sciences of the United States of America 110(27): 11169–11174.

Slifstein, M. and M. Laruelle (2001). “Models and methods for derivation of in vivo neuroreceptor parameters with PET and SPECT reversible radiotracers.” Nucl Med Biol 28(5): 595–608.

Small, D. M., M. Jones-Gotman and A. Dagher (2003). “Feeding-induced dopamine release in dorsal striatum correlates with meal pleasantness ratings in healthy human volunteers.” NeuroImage 19(4): 1709–1715.

Small, D. M., M. G. Veldhuizen, J. Felsted, Y. E. Mak and F. McGlone (2008). “Separable Substrates for Anticipatory and Consummatory Food Chemosensation.” Neuron 57(5): 786–797.

Smith, K. S., A. J. Tindell, J. W. Aldridge and K. Berridge (2009). “Ventral Pallidum Roles in Reward and Motivation.” Behavioural Brain Research 196(2): 155–167.

Soares-Cunha, C. and J. A. Heinsbroek (2023). “Ventral pallidal regulation of motivated behaviors and reinforcement.” Front Neural Circuits 17: 1086053.

Stunkard, A. J. and S. Messick (1985). “The three-factor eating questionnaire to measure dietary restraint, disinhibition and hunger.” Journal of Psychosomatic Research 29(1): 71–83.

Sun, X., M. G. Veldhuizen, A. E. Wray, I. E. de Araujo, R. S. Sherwin, R. Sinha and D. M. Small (2014). “The neural signature of satiation is associated with ghrelin response and triglyceride metabolism.” Physiology & Behavior 136: 63–73.

Tataranni, P. A., J. F. Gautier, K. Chen, A. Uecker, D. Bandy, A. D. Salbe, R. E. Pratley, M. Lawson, E. M. Reiman and E. Ravussin (1999). “Neuroanatomical correlates of hunger and satiation in humans using positron emission tomography.” Proc Natl Acad Sci U S A 96(8): 4569–4574.

Tellez, L. A., X. Ren, W. Han, S. Medina, J. G. Ferreira, C. W. Yeckel and I. E. de Araujo (2013). “Glucose utilization rates regulate intake levels of artificial sweeteners.” The Journal of physiology 591(22): 5727–5744.

Thanarajah, S. E., H. Backes, A. G. DiFeliceantonio, K. Albus, A. L. Cremer, R. Hanssen, R. N. Lippert, O. A. Cornely, D. M. Small, J. C. Brüning and M. Tittgemeyer (2018). “Food Intake Recruits Orosensory and Post-ingestive Dopaminergic Circuits to Affect Eating Desire in Humans.” Cell metabolism 0(0).

van der Plasse, G., R. van Zessen, M. C. Luijendijk, H. Erkan, G. D. Stuber, G. M. Ramakers and R. A. Adan (2015). “Modulation of cue-induced firing of ventral tegmental area dopamine neurons by leptin and ghrelin.” Int J Obes (Lond) 39(12): 1742–1749.

van Galen, K. A., A. Schrantee, K. W. ter Horst, S. E. la Fleur, J. Booij, R. T. Constable, G. J. Schwartz, R. J. DiLeone and M. J. Serlie (2023). “Brain responses to nutrients are severely impaired and not reversed by weight loss in humans with obesity: a randomized crossover study.” Nature Metabolism 5(6): 1059–1072.

van Vliet, S., H. E. Koh, B. W. Patterson, M. Yoshino, R. LaForest, R. J. Gropler, S. Klein and B. Mittendorfer (2020). “Obesity Is Associated With Increased Basal and Postprandial β-Cell Insulin Secretion Even in the Absence of Insulin Resistance.” Diabetes 69(10): 2112–2119.

Vickery, T. J., M. M. Chun and D. Lee (2011). “Ubiquity and specificity of reinforcement signals throughout the human brain.” Neuron 72(1): 166–177.

Wang, G.-J., D. Tomasi, A. Convit, J. Logan, C. T. Wong, E. Shumay, J. S. Fowler and N. D. Volkow (2014). “BMI Modulates Calorie-Dependent Dopamine Changes in Accumbens from Glucose Intake.” PLoS ONE 9(7): e101585–e101585.

Weber, L. A., D. M. Yee, D. M. Small and F. H. Petzschner (2025). “The interoceptive origin of reinforcement learning.” Trends in Cognitive Sciences 29(9): 840–854.

Weir, J. B. d. V. (1949). “New methods for calculating metabolic rate with special reference to protein metabolism.” J Physiol 109(1-2): 1–9.

Wilson, S. M., A. P. Maes, C. J. Yeoman, S. T. Walk and M. P. Miles (2021). “Determinants of the postprandial triglyceride response to a high-fat meal in healthy overweight and obese adults.” Lipids Health Dis 20(1): 107.

Yousif, N., R. Z. Fu, B. Abou-El-Ela Bourquin, V. Bhrugubanda, S. R. Schultz and B. M. Seemungal (2016). “Dopamine Activation Preserves Visual Motion Perception Despite Noise Interference of Human V5/MT.” The Journal of Neuroscience 36(36): 9303–9312.

